# A long-lived human gastric tissue model defines regulators of lineage maturation

**DOI:** 10.64898/2026.11.25.769210

**Authors:** Ninouk Akkerman, Mikhail Nikolaev, Lukas Adam, Yannik Bollen, Jannika Bosch, Suraj Rajan Vasandani, Laura Gaspa-Toneu, Andrea Corsinotti, Alberto Valdeolivas, Franziska Voellmy, Irineja Cubela, Rebecca A. Siwicki, Javier Torres-Barajas, Hans Clevers, J. Gray Camp, Nikolche Gjorevski, Joep Beumer

## Abstract

The gastric epithelium maintains barrier function against food-borne pathogens and executes metabolic activities through specialized cell types. The niche signals and gene-regulatory networks governing gastric specification remain poorly understood, partly due to the lack of mature human in vitro models. Here we develop long-lived engineered gastric epithelia that recapitulate gland-pit topography and, upon extended culture, give rise to all known gastric epithelial lineages including rare enteroendocrine and tuft cell subtypes. We use niche and genetic perturbations on EGE to explore patterning, cell differentiation, and gene regulation in the stomach. We establish a role for Neuregulin-1 in foveolar cell maturation, discover a novel FOXI1-regulated population resembling ionocytes and generate a roadmap of gastric enteroendocrine cell differentiation. This work establishes a physiological model that provides insights into the regulation of lineage maturation and can be employed to investigate broad aspects of gastric homeostasis and disease.

## Main

The stomach plays an integral role in digestion. Three anatomical regions (cardia, corpus, and antrum) contain numerous cell types that aid in digestion and protect against a hostile luminal environment. The simple columnar epithelium is organized in flask-shaped glands, which feed into individual pit regions^1–3^. The pit is composed of mucus-producing foveolar cells providing a barrier against the acidic lumen and food-borne pathogens. Corpus glands harbor the majority of proton-pumping parietal cells, digestive enzyme-secreting chief cells, immune-regulatory tuft cells, as well as multiple enteroendocrine cells (EEC) subtypes orchestrating physiological processes including acid secretion and hunger^4,5^ (Extended Data Fig. 1a).

In contrast to the adjacent intestine, the stem cell hierarchy in the gastric epithelium is ill-defined^1^. This is partly due to the lack of promoter-based tools that label gastric stem cells specifically, with the exception of the recently discovered AQP5+ cells in the antrum^6,7^. Additionally, tamoxifen treatment depletes parietal cells and causes chief cell metaplasia, compromising the study of homeostatic cellular behaviour^8–10^. As a result, the niche factors and downstream gene-regulatory networks that govern gastric fates remain largely unknown. While gastric organoids derived from adult stem cells (ASCs) and induced pluripotent stem cells (iPSCs) promise to address these questions, significant challenges remain. ASC-derived organoids display limited cellular diversity and lack physiological patterning^11,12^. iPSC-derived cultures include epithelial, mesoderm and ectoderm cell types, enabling studies of tissue-level interactions. However, they model developmental stages and require xenografting into immunocompromised rodent hosts to promote maturation^13–15^.

We combined ASC-derived organoids, ECM scaffold- and device microengineering to establish long-lived human gastric tissues that feature a reproducible gland-pit epithelial architecture. We show that the gland-pit geometry confers *in vivo*-like cell type patterning, whilst long-term growth facilitates the emergence of all epithelial lineages. The model captures all known gastric EEC subtypes, including rare PYY- and GLP1-producing L cells. Additionally, we identified a previously unreported gastric cell type reminiscent of ionocytes, which regulate ion balance in other epithelial tissues^16–21^. Coupling spatial analyses of primary tissues with functional studies in EGE, we defined regulators of gastric lineage maturation, finding that Neuregulin 1 (NRG1) promotes foveolar cell maturation. Moreover, we identified FOXI1 as a driver of ionocyte differentiation, and established a role of ASCL1 in gastric enterochromaffin cell differentiation. Thus, EGE enable previously unattainable developmental and functional interrogation of human gastric cell types. Capturing all key lineages of the epithelium, we envision that this model will aid studies of human stomach homeostasis and disease.

### Topographical and temporal cues drive gastric lineage maturation

Lack of cell-type diversity and patterning are well-known deficiencies of ASC-derived gastric organoids^22^. We confirmed this limited differentiation by establishing a quintuple reporter (5R) organoid line^23^ that fluorescently labels markers for Ghrelin-(GHRL-) producing EECs (X cells), chief cells (Lipase F, LIPF), parietal cells (ATPase H+/K+ Transporting Subunit Alpha, ATP4A), foveolar cells (Gastrokine 1, GKN1) and nuclei (Histone-2B1C, H2BC4) (Extended Data Fig. 1b-g). 5R organoids, which generally require fortnightly passaging^11^, were cultured and imaged over the course of 4 weeks. As demonstrated previously, the organoids lacked mature lineages and spatial patterning (Fig. 1a), which we attribute to the absence of symmetry-breaking cues and short culture times incongruent with the slower turnover of mature gastric epithelial cells^24-26^.

**Fig. 1:**
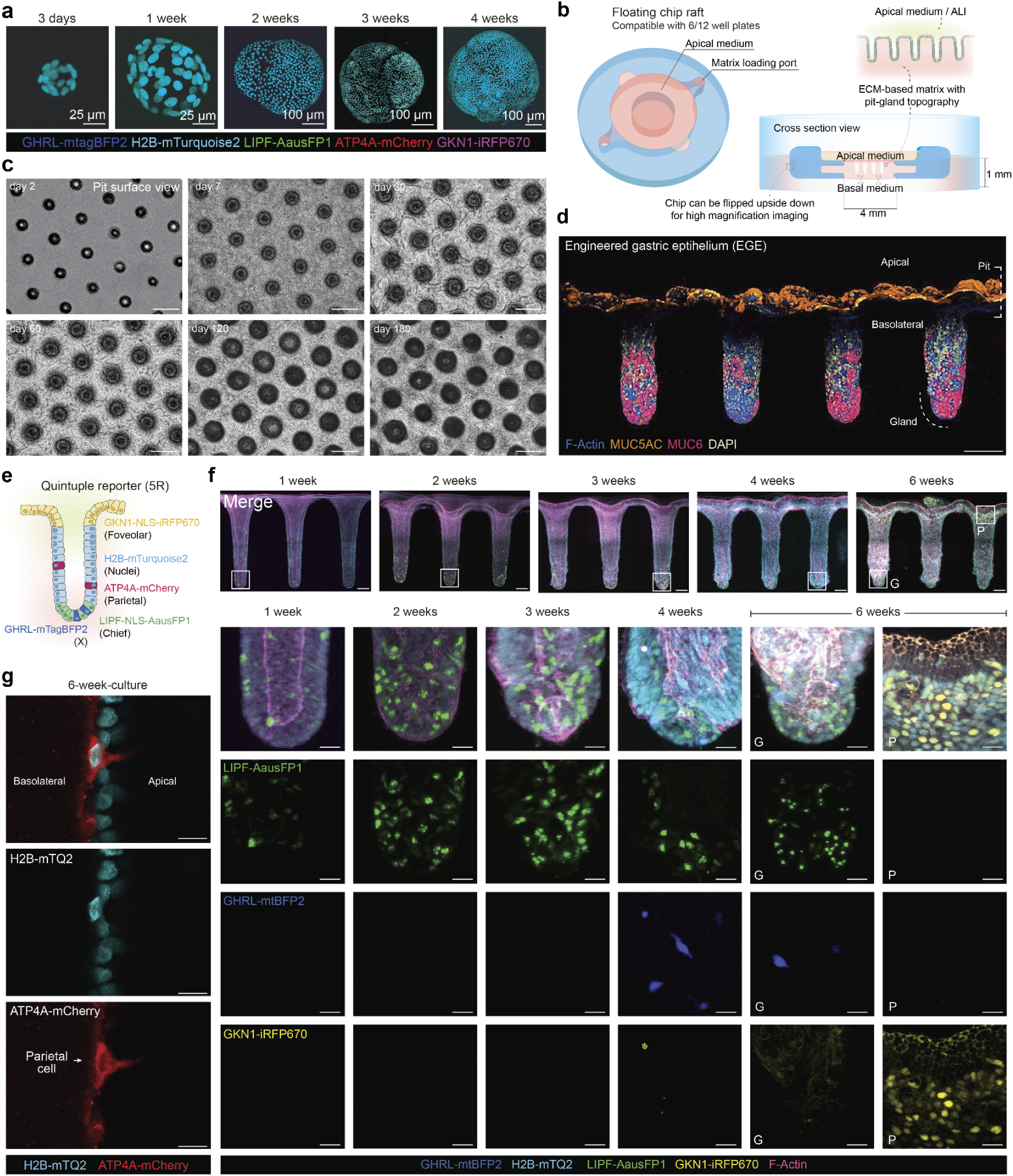
Homeostatic, patterned corpus tissues through long-term culture on engineered scaffolds. **a**, Representative fluorescent images of stomach organoids with H2B, GKN1, ATP4A, LIPF and GHRL reporters at day 3 and at 1, 2, 3, and 4 weeks of culture. **b**, Schematic representation of engineered stomach tissue culture shown in top-down view (left) and cross section (right), demonstrating recapitulation of pit–gland architecture generated by microstamping. **c**, Multiplex immunofluorescence image of engineered corpus epithelium at 5 weeks of culture, showing MUC5AC and MUC6. Scale bar: 100 μm. **d**, Diagram of EGE patterning. **e,** ATP4A–mCherry reporter signal highlighting a parietal cell within the engineered epithelium. Scale bar: 10 μm. **f**, Bright-field images of engineered stomach tissues at days 2, 7, 60, 120, and 180 of culture. Scale bars: 200 μm. **g**, Multiplex fluorescence imaging of engineered corpus epithelia at 1, 2, 3, 4, and 6 weeks of culture, showing GKN1, GHRL, ATP4A, LIPF, H2B reporters and F-actin staining. Scale bars overview: 100 μm, inserts: 25 μm. G: gland, P: pit.

We previously demonstrated that introducing crypt-like domains is sufficient to induce and control symmetry breaking of intestinal organoids^27–30^. Building on this concept, we used microstamping technology to impart a corpus gland-pit topography on the surface of ECM-based hydrogels (Fig. 1b, Extended Data Fig. 2a). To extend the lifespan of the cultures, we developed a new culture micro-device, whose floating design facilitates nutrient and growth factor diffusion, gas exchange, and waste removal (Fig. 1b). Seeding single cells derived from corpus organoids on the hydrogel yielded confluent, apically accessible epithelia that adopted the surface topography, forming reproducible gland-pit domains (Fig. 1c). Remarkably, we found that the topographically structured corpus epithelia cultured in the device could be stably grown for at least 6 months without passaging (Fig. 1c, Supplementary Video 1). We termed this model ‘engineered gastric epithelium’ (EGE).

Staining the EGE for MUC5AC and MUC6 after 5 weeks of culture revealed striking patterning of the two mucins, with MUC5AC (labeling foveolar cells) localizing to the surface and MUC6 (labeling neck cells) below (Fig. 1d). Proliferation was likewise patterned; it occurred throughout the gland but was virtually absent from the differentiated pit surface (Extended Data Fig. 2b).

To assess the emergence of other mature lineages, we seeded 5R organoids and imaged the engineered epithelia weekly for 6 weeks (Fig. 1e-f). Chief cells appeared after 1 week of culture and became localized to the bottom of the glands (Fig. 1e, Supplementary Video 2). From week 4 onwards, X cells were visible at the gland base and at 6 weeks of culture the surface epithelium was fully populated by foveolar cells (Fig. 1f). Additionally, we observed rare parietal cells in the glands of 6-week old cultures (Fig. 1g, Supplementary Video 3). To further investigate the differentiation dynamics, we performed scRNAseq of EGEs at 4 time points (week 2, 3, 4 and 6). We identified all major cell types, apart from parietal cells (Fig. 2a-b, Extended Data Fig. 2c), likely owing to their relative rarity or fragility. Flow cytometry indeed suggested low abundance (mean = 3.0% ± 0.23 SEM, 10 independent EGEs) of ATP4A-mCherry+ cells in 6-week old cultures (Fig. 2c, Supplementary Fig. 1a), yet qPCR analysis revealed a >60-fold increase in expression of ATP4A and 5-fold upregulation of ATP4B in mCherry+ compared to mCherry-cells (Fig. 2d, 6 independent EGE). Other rare lineages such as chief, tuft and enteroendocrine cells increased in proportion from 2 to 6 weeks of culture (Fig. 2e). These data suggest that prolonged culture is crucial for the differentiation and maturation of corpus lineages, with certain cell types requiring as long as 6 weeks of uninterrupted culture.

**Fig. 2:**
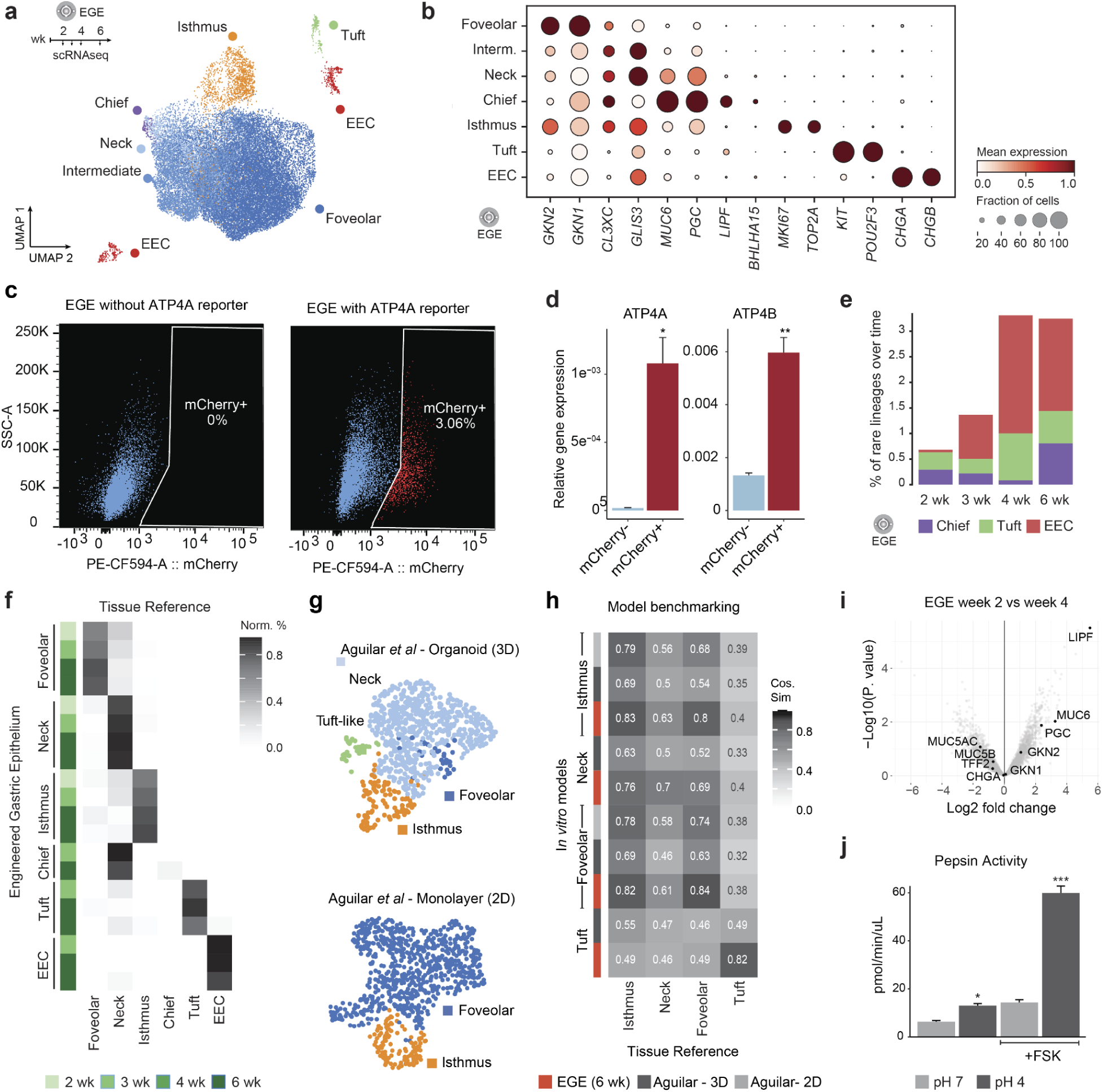
Mature epithelial lineages arise upon long-term EGE culture. **a**, Integrated single-cell RNA-seq atlas of corpus cultures with annotated epithelial cell types. **b**, Dot plot showing expression of canonical marker genes used for cell-type annotation. **c**, Flow cytometry analysis (side scatter area (SSC-A) versus mCherry) of engineered corpus tissue (EGE) carrying an ATP4A–mCherry reporter and controls without reporter; ATP4A–mCherry–positive cells are detected at approximately 3% in corpus reporter cultures (N=10 separate EGE). **d**, Barplot shows relative expression levels of *ATP4A* and *ATP4B* relative to *GAPDH* and *ACTB* in mCherry-positive and mCherry-negative cell populations (qPCR analysis of N=6 replicate EGE, FAC-sorted for mCherry-positive and mCherry-negative populations). **e**, Temporal dynamics of rare epithelial lineage proportions over culture duration, as detected by scRNAseq. **f**, Comparison of normalized predicted cell-type annotations based on a reference tissue atlas with experimentally annotated cell types. **g**, UMAP embedding of reference single-cell RNA-seq datasets from Aguilar *et al.* (3D gastric organoids and 2D stomach monolayers) with annotated epithelial cell types. **h**, Heatmap showing maximum cosine similarity of gene expression profiles between EGE, Aguilar 2D monolayers, Aguilar 3D organoids, and adult human stomach tissue reference. **i**, Volcano plot of differential protein abundance secreted in the apical supernatant between 2-week and 4-week-old EGE. **j**, Apical pepsin activity measurements in control and forskolin (FSK)-treated cultures across two pH levels.

Indeed, benchmarking EGE against a reference atlas constructed from adult stomach scRNAseq datasets^31,32^ confirmed that EGE-derived gastric cell types most closely resemble their primary tissue counterparts after 3 or more weeks of culture (Fig. 2f). In contrast, short-lived conventional corpus organoids and monolayers^22^ showed reduced cellular diversity and similarity to tissue (Extended Data Fig. 2d). Organoids were composed predominantly of neck mucous and isthmus cells, with occasional foveolar cells and tuft cell progenitors (Fig. 2g). Monolayers exclusively contained foveolar and isthmus cells. Although a separate neck/chief cell population was previously annotated in organoids^22^, we could not identify hallmarks of mature chief cells, including co-expression of pepsinogens and lipases (encoded by LIPF). Though EGE chief cells displayed reduced tissue resemblance score than other EGE lineages and did not express type A pepsinogens (PGA3, PGA5, Fig. 2f), they robustly expressed lipases, as well MIST1^33^ (encoded by BHLHA15, Fig. 2b) – a principal regulator of chief cell identity.

We evaluated the functional maturity of the EGE model by using mass spectrometry to analyze the presence of proteins involved in digestive and endocrine processes. We detected secretion of mucins (MUC5AC, MUC5B, MUC6), trefoil factors (TFF2, TFF3), gastrokines (GKN1, GKN2) and digestive enzymes (PGC, LIPF). The abundance of digestive enzymes increased at week 4 of culture compared to week 2, in line with the maturation of the cultures observed at the RNA level (Fig. 2i, Extended Data Fig. 2e, Supplementary Table 1). Furthermore, we found that EGE produce functional digestive enzymes, as evidenced by apical pepsin activity, which, as expected, was positively regulated by an acidic environment (pH 4) and further amplified upon stimulation with a cyclic adenosine monophosphate (cAMP) agonist^34–36^ (Fig. 2j).

Altogether, our data implicate time as a critical determinant of gastric cell maturation. By establishing long-term, passage-free cultures of corpus epithelium, we have uniquely enabled the appearance of all epithelial lineages in a single medium condition. Furthermore, our bioengineering approach enabled the formation of apically accessible corpus epithelia with gland-pit domains, conferring reproductive cell-type patterning.

### Spatial transcriptomics reveals patterned niche factors controlling gastric cell differentiation

Differentiation and cell type distribution of the gastric epithelium is guided by niche factors that are differentially concentrated along the gland-pit axis. To map the spatial distribution of ligands and receptors in stomach tissue, we performed spatial transcriptomics (Xenium) on corpus and antrum tissue (2 donors, matched) using a probe-based approach with subcellular resolution37. We curated a probe set of 480 genes marking the different epithelial, mesenchymal and immune populations, as well as ligands and receptors identified in gastric tissue^4,31,32^ (Supplementary Table 2). Immunofluorescence-based cell segmentation resolved all major cell classes (Fig. 3a-e, Extended Data Fig. 3a-c, Supplementary Table 3).

**Fig. 3:**
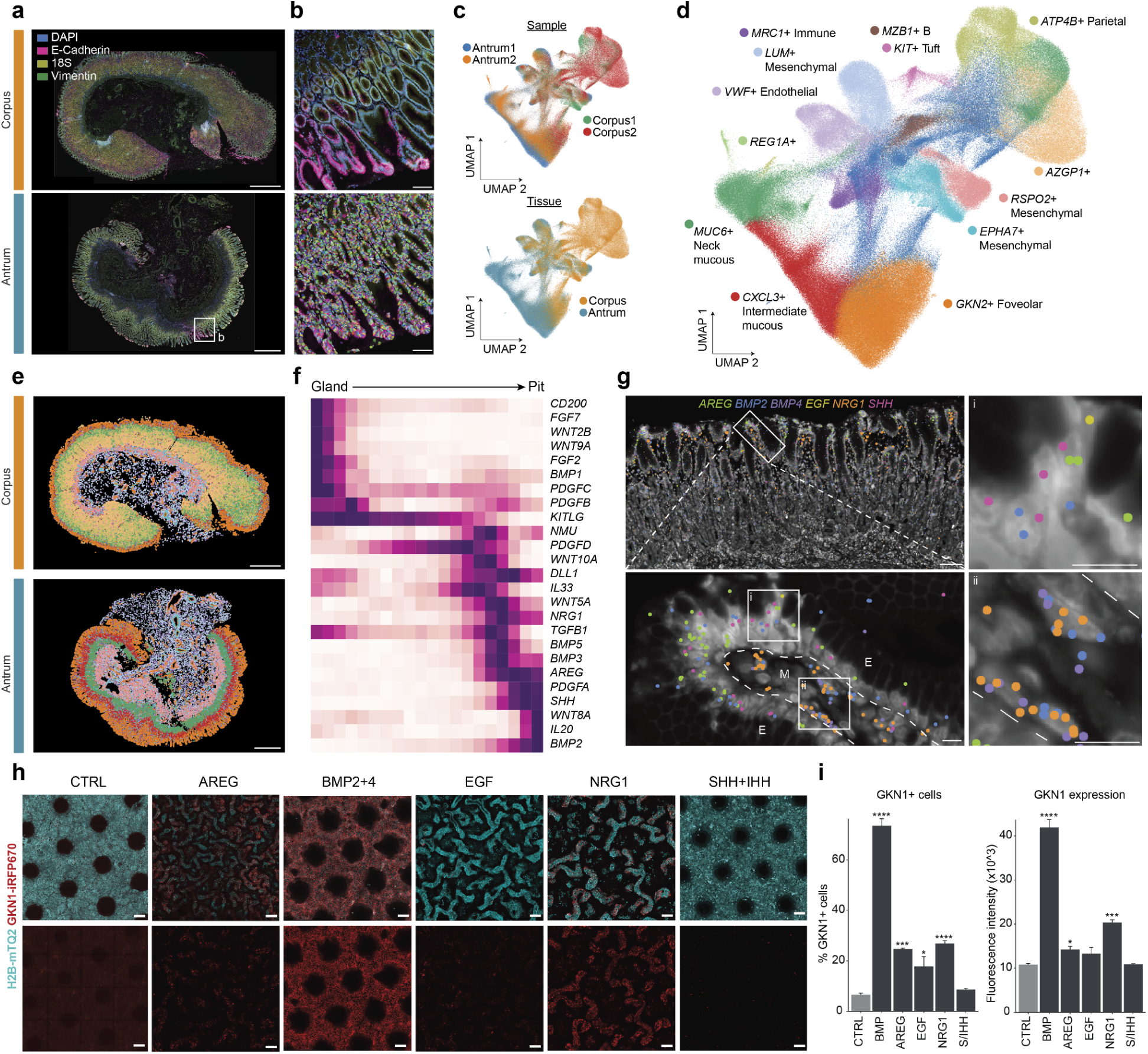
Spatial transcriptomics reveals patterned ligands driving foveolar differentiation. **a**, Representative Xenium spatial transcriptomics images from two corpus and two antrum tissue samples, showing merged channels for DAPI, E-cadherin, 18S rRNA, and vimentin. Corpus/Antrum1 and 2 correspond to Donors 2 and 3, respectively. Scale bars: 1 mm. **b**, Higher-magnification view highlighting gastric pit–gland architecture and single-cell segmentation within the Xenium dataset showing selected marker expression (top) and cell segmentation (bottom). Scale bars: 100 μm. **c**, UMAP embedding derived from Xenium gene expression profiles, colored by individual sample (top) and anatomical region (bottom). **d**, UMAP embedding with annotated epithelial and non-epithelial cell types identified from spatial transcriptome data. **e**, Spatial projection of cell-type annotations onto the tissue architecture shown in **a**, with segmented cells colored by cell type. **f**, Spatial enrichment analysis of selected ligand transcripts along the gland-to-pit axis in corpus tissue. **g**, Representative views showing spatial expression patterns of *AREG*, *BMP2*, *BMP4*, *EGF*, *NRG1*, and *SHH* in corpus tissue (Donor 2). Each dot represents a single transcript. Scale bar overview: 100 μm, inserts: 10 μm. E: epithelium, M: mesenchyme. Dotted line: boundary between epithelium and mesenchyme. **h**, Representative fluorescence images of 4-week-old engineered stomach cultures following treatment with indicated ligands for 2 weeks, showing H2B and GKN1 reporters. Top-down view. Scale bars: 100 μm. **i**, Quantitative flow cytometry analysis of GKN1-iRFP670+ cells in whole 4-week-old cultures treated for 2 weeks with the indicated growth factors (n=3 biological replicates per condition), showing the percentage of GKN1-iRFP670+ cells and the relative fluorescent intensity of the positive population.

To quantify the spatial distribution of niche factors, we generated epithelial masks for each tissue, ordered cells from gland bottom to pit, and projected ligand expression onto a two-dimensional spatial gradient (Fig. 3f, Extended Data Fig. 3d). In line with their reported role in foveolar cell differentiation in rodents, we found preferential localization of *BMP2* and *SHH* in the pit^38–40^ (Fig. 3f, Supplementary Table 4). This *SHH* expression pattern contrasts with earlier findings that described parietal cells as the predominant source^41^. However, scRNAseq data confirmed *SHH* expression in foveolar cells, without significant expression in parietal cells (Extended Data Fig. 3e).

While EGF signaling is reportedly involved in foveolar cell maturation^38^, we did not find high *EGF* expression in the pit zone. Instead, there was abundant expression of other ErbB ligands, including *AREG* and *NRG1* (Fig. 3f). *SHH*, *AREG* and *EGF* were expressed in the foveolar epithelium, while *BMP4* and *NRG1* were expressed by stromal cells beneath (Fig. 3g, Extended Data Fig. 3e). *BMP2* was expressed both in foveolar cells, as well as stromal cells in the pit vicinity.

EGE, which capture all epithelial lineages and are amenable to experimental manipulation, allow us to query the functional roles of putative regulators identified by the spatial transcriptomic analysis. We supplemented 5R EGE with these ligands (AREG, EGF, NRG1 and SHH) as well as BMPs for 2 weeks, and explored their effects on epithelial differentiation. We could recapitulate the reported effect of BMP signaling in rodents^40^, evidenced by a large expansion of the number of GKN1+ cells as well as an increase in GKN1 expression levels (Fig. 3h-i, Supplementary Fig. 1b). In line with their localization, we found that AREG and NRG1 caused a substantial increase in the number of foveolar cells that was higher when compared with EGF. Only NRG1 induced higher levels of GKN1-iRFP670 signal per cell (Fig. 3i), suggesting that these ligands stimulate pit cell expansion and maturation to varying degrees. SHH and IHH, previously described to induce foveolar differentiation^42^, did not affect GKN1+ cell numbers or levels, as measured by flow cytometry (Fig. 3h-i). Foveolar cells produce significant amounts of Hedgehog ligands, and this may explain why exogenous stimulation does not further enhance their differentiation (Extended Data Fig. 3e). Additionally, it is possible that foveolar cell-derived SHH does not act in an autocrine manner, but rather instructs stromal cell production of BMPs, as has been demonstrated in intestinal villi^43,44^.

To profile the impact of niche factors on corpus differentiation more comprehensively, we exposed EGE to various ligands (21 conditions, n=3) and performed bulk RNA sequencing. This screen confirmed the role of BMPs and NRG1 in enhancing foveolar differentiation (Extended Data Fig. 3f-g, Supplementary Table 5). Additionally, we found that LATS inhibition (activating YAP/TEAD signaling) strongly induced chief cells, while ErbB activation caused a decline in the EEC population (Extended Data Fig. 3g). The latter has been reported in the intestine^45^. Together, our experiments establish the zonation and functional role of human gastric morphogens.

### Identification of FOXI1-regulated gastric ionocytes

EGE capture a significant (0.5-1%) population of mature gastric tuft cells (Fig. 2a, 4a) that have been largely inaccessible due to their rarity and lack of *in vitro* tools. Tuft cells are chemosensory cells that have been described to be essential for adaptation to dietary chitin, inducing gastric production of digestive enzymes and improving metabolic health^46^. Extensive scRNAseq analyses of intestinal tuft cells have revealed cellular heterogeneity, including 4 separate subtypes, but their heterogeneity has not been explored in the stomach^47–49^. We therefore subclustered the EGE tuft cells and resolved two distinct cell populations. One expressed ‘classical’ tuft cell markers including high *KIT*, *POU2F3*, *BMX* and *AVIL*. Strikingly, the second population of *KIT+*/*POU2F3+* cells expressed markers reminiscent of ionocytes previously described in the lung^16,17^, such as *FOXI1*, *ASCL3* and *TFCP2L1* (Fig. 4b-d). These cells were negative for *BMX*, *ALOX5* and *AVIL,* disqualifying them as tuft cells. We could also distinguish these separate cell populations *in vivo* based on our spatial transcriptomic data (Fig. 4e).

**Fig. 4:**
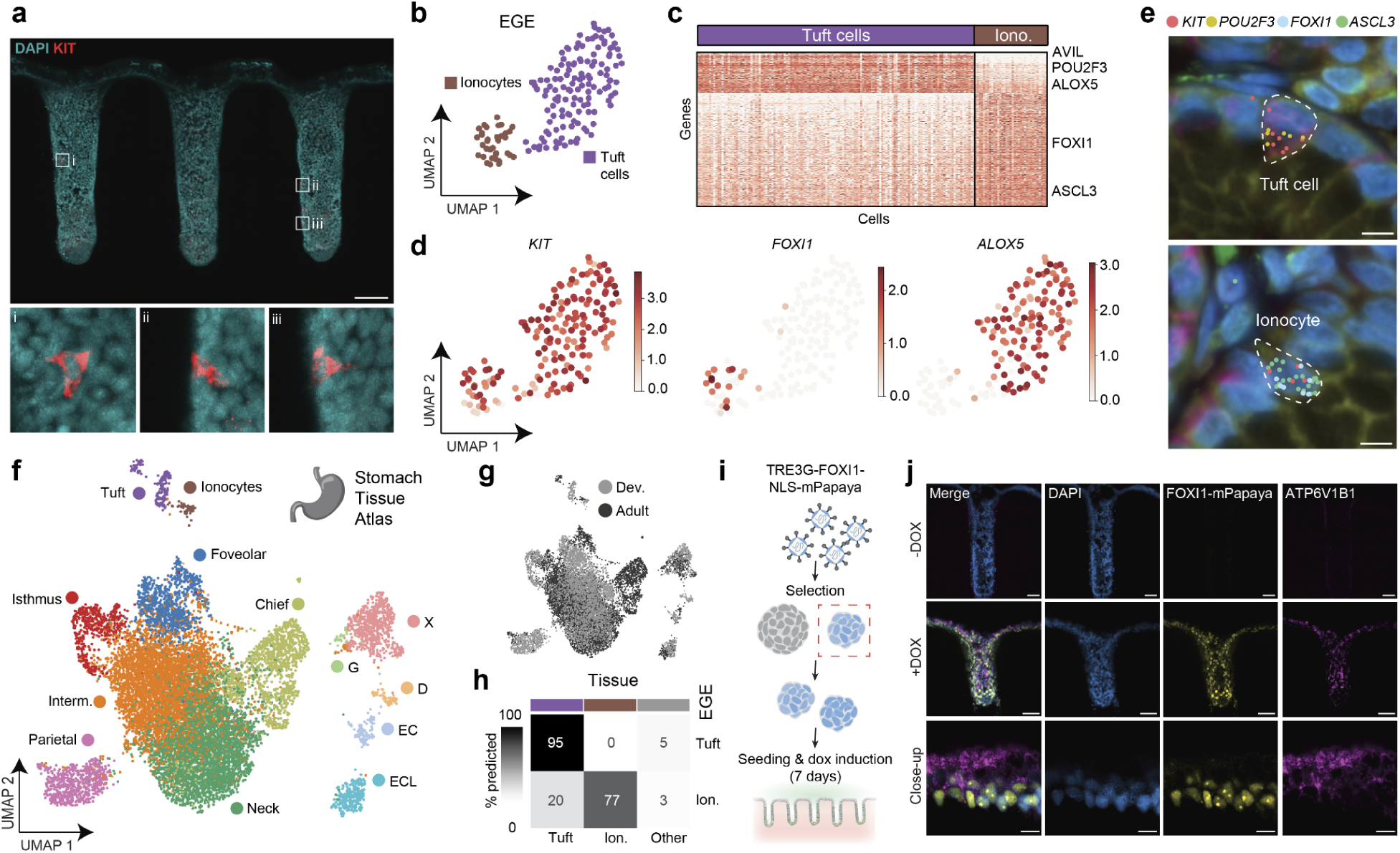
Gastric tissue harbors *FOXI1*-regulated ionocytes. **a**, Immunofluorescence imaging of 4-week-old EGE stained for KIT and DAPI. Scale bars: 100 μm. **b**, UMAP embedding of sub clustered EGE-derived tuft cells, revealing a second cluster resembling ionocytes, annotated by cell type. **c**, Differential gene expression between EGE tuft cells and ionocytes, showing key markers enriched in either cluster (*AVIL*, *POU2F3* and *ALOX5* in tuft cells, *FOXI1* and *ASCL3* in ionocytes). **d**, UMAP embedding of EGE tuft cells and ionocytes, showing expression of *KIT*, *FOXI1* and *ALOX5*. **e**, Immunofluorescent images derived from the Xenium analysis of adult corpus tissue, showing a tuft cell with transcripts for *KIT* and *POU2F3* and an ionocyte with transcripts for *KIT*, *FOXI1* and *ASCL3*. Each dot represents one transcript. Scale bar: 5 μm. **f**, UMAP embedding of the integrated single cell RNAseq tissue atlas annotated by cell type. **g**, UMAP embedding of the integrated single cell RNAseq tissue atlas annotated by tissue source, with cells derived from the developing corpus in gray and those derived from adult tissue in black. **h**, Correlation plot showing resemblance score (value between 0 and 100) of EGE-derived tuft cells and ionocytes to those derived from the integrated atlas. **i**, Diagram showing the workflow of the overexpression experiment. **j**, Immunofluorescence imaging of EGE upon 7-day overexpression of FOXI1-mPapaya1 versus control, showing DAPI, mPapaya1 and ATP6V1B1. Scale bars top and middle: 100 μm, bottom: 10 μm.

Next, we sought to benchmark EGE ionocytes to their tissue counterparts as well as compare them to other described ionocyte-like cell populations. To resolve their presence and signature across gastric development and adulthood, we generated a scRNAseq dataset of the developing human stomach while enriching for rare lineages via FAC-sorting (c-KIT for tuft cells^48^ and ionocytes; CD200 for EECs^50^) and integrated these data with our adult tissue reference (Fig. 4f, Extended Data Fig. 4a-b). We found ionocytes in both developing and adult stomach tissue (Fig. 4f-g). EGE tuft cells and ionocytes displayed high concordance to their tissue counterparts (Fig. 4h). We correlated gastric ionocyte signatures to a previously published integrated cross-tissue ionocyte atlas^51^, comprising ionocytes from thymus^52^, epididymis^53^, lung^16,54,55^ and skin^52,56^ and kidney intercalated cells^57^. Out of these tissues, gastric ionocytes most strongly resembled pulmonary ionocytes, followed by epididymis and thymus (Extended Data Fig. 4c). Several genes - such as *SOX4*, *ATP6V0D2*, *SEMA3C*, *AMACR*, and *ASCL3* - were shared between pulmonary and gastric ionocytes, but were absent from kidney intercalated cells, skin, epididymis, and thymus (Extended Data Fig. 4d).

Differentiation of pulmonary ionocytes is mediated through the transcription factor *FOXI1*, while tuft cells across epithelial tissues are regulated by various POU-domain-containing transcription factors and co-activators (including *POU2F3*^58^ and *POU2AF2*^59^). To test the impact of these TFs on gastric differentiation, we transiently overexpressed *FOXI1* and *POU2AF2* in EGE. Doxycycline-mediated induction of FOXI1 for 7 days led to a significant increase in the expression of ionocyte-associated genes (Fig. 4i-j, Extended Data Fig. 4e). Tuft cell markers on the other hand were largely unaffected. In contrast, overexpression of POU2AF2 induced canonical tuft cell markers while having limited effect on ionocytes. This indicates that - even though POU2F3/POU2AF2 are expressed in ionocytes - this module mainly drives tuft cell differentiation. Instead, FOXI1 suffices to induce ionocyte but not tuft cell differentiation.

### EGE contain mature gastric enteroendocrine cells

EGE capture EEC subtypes that were not generated in previous adult models, even through cell fate engineering^50^: All subtypes including X, D (producing somatostatin), enterochromaffin-like (ECL, producing histamine) and enterochromaffin (EC, producing serotonin) cells were identified via immunostaining of their main hormone product, as well as in EGE scRNAseq data (Fig. 5 a-c, Supplementary Video 4). To assess functional maturity of the different EEC subtypes captured in EGE, we stimulated hormone secretion through cAMP agonism in EGE. We measured increased hormone secretion over culture time, suggesting progressive maturation or increase of EEC numbers (Fig. 5c).

**Fig. 5:**
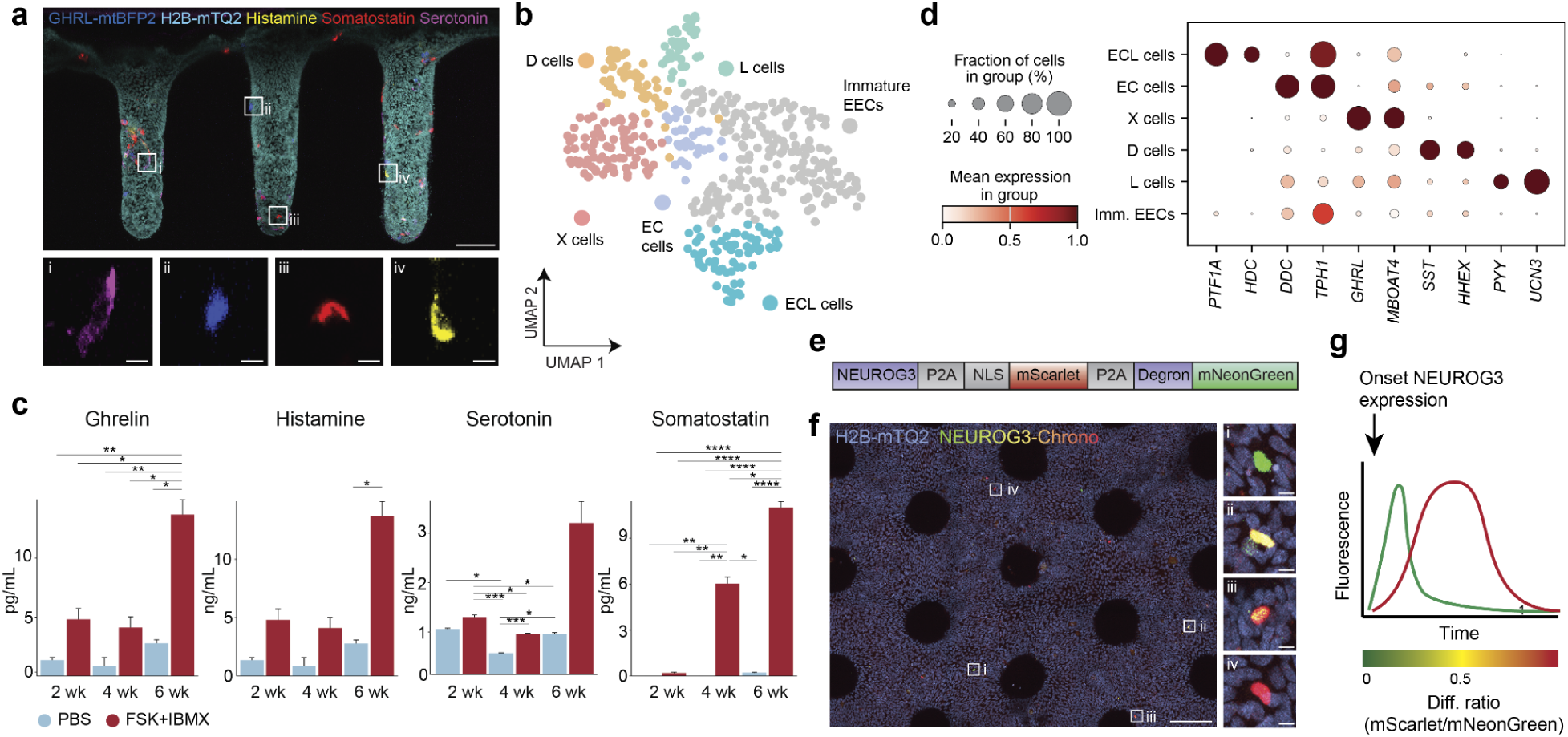
EGE include all gastric corpus enteroendocrine cell types. **a**, Multiplexed fluorescent images of 4-week-old EGE showing hormone expression by fluorescent reporter (GHRL-P2A-mTagBFP2) and immunostaining (Serotonin (magenta), Somatostatin (red), Histamine (yellow)). Scale bar overview: 100 μm, inserts: 10 μm. **b**, UMAP showing populations of EEC subtypes derived from EGE. **c**, Dot plot showing the mean expression of EEC subtype-enriched genes in EECs derived from EGE. **d**, Concentrations of corpus hormones Ghrelin, Histamine, Serotonin and Somatostatin in the apical supernatant after 6 hours in PBS (blue) or stimulated with Forskolin and IBMX (FSK+IBMX, red). **e**, Construct encoding the NEUROG3-Chrono reporter. **f**, IF images showing NEUROG3-Chrono expressing cells in EGE. Scale bar overview: 100 μm, inserts: 5 μm. **g**, Fluorescence dynamics of the NEUROG3-Chrono reporter. Right: Diagram showing the NEUROG3-Chrono reporter fluorescence dynamics.

Additionally, EEC subclustering resolved rare L-like cells expressing *GCG* and *PYY*, which also produce high levels of *UCN3* coding for Urocortin 3 (Fig. 5d). This peptide is co-released with glucagon and insulin in pancreatic endocrine cells and provides negative feedback on hormone secretion by inducing somatostatin release^60^. L cells are exceedingly rare in the stomach but have been reported to increase upon bariatric surgery^61^. L cells *in vitro* predominantly express Prohormone Convertase Subtilisin/Kexin type 1 (*PCSK1*) and to a lesser extent *PCSK2* (Extended Data Fig. 5a). Both enzymes cleave the prohormone proglucagon into active peptide hormones, with *PCSK1* producing (among other peptides) GLP-1 and *PCSK2* generating glucagon^62^. IF staining confirmed the presence of GLP1+ cells in human corpus tissue (Extended Data Fig. 5b). Given their paucity, it is plausible that these cells have paracrine functions rather than impacting systemic glucose homeostasis. We therefore assessed the expression of GLP-1 receptors (encoded by *GLP1R)* in tissue and found predominant production in the endocrine lineages - particularly in X cells (Extended Data Fig. 5c). In line with this, we could verify *GLP1R* expression in Ghrelin-producing X cells in the Xenium dataset (Extended Data Fig. 5d) - suggesting locally produced GLP-1 may affect secretion of this hunger hormone.

### A regulatory roadmap of gastric enteroendocrine differentiation

While fate regulators of various EEC subtypes are described in the intestine, there is currently little known about determinants of their differentiation in the gastric epithelium. To explore differentiation of human gastric EECs, we exploited the fact these are naturally produced in EGE. Given that EEC turnover is infrequent and progenitor states are thus rare, we employed a modified version of a previously developed dual-fluorescent reporter termed NEUROG3-Chrono^64^ (Fig. 5e, Extended Data Fig. 1h). A fast-folding FP (mNeonGreen) is destabilized by a degron element and separated by a P2A from a slow-folding FP with a slow turnover (mScarlet). This approach exploits the transient nature of *NEUROG3* expression reported in other tissues. EEC progenitors consequently express a short pulse of mNeonGreen followed by a prolonged pulse of mScarlet (Fig. 5f). The ratio of mNeonGreen/mScarlet fluorescence intensity (differentiation ratio) reflects a temporal differentiation axis.

NEUROG3-Chrono+ cells in EGE represented a rare population (Fig. 5g). Obtaining sufficient numbers of EEC progenitors for single-cell transcriptomic profiling required substantially (∼40x) larger epithelia. We thus extended our earlier manufacturing process to build a new floating chip-carrier for EGE, supporting ∼2×2 cm engineered epithelia (Fig. 6a-b, Supplementary Video 5). 10 million reporter cells were seeded per macroscale chip and harvested 2-3 weeks post-seeding. We performed an index FAC-sort of EEC progenitors at different maturation stages (Fig. 6c-d, Supplementary Fig. 1c) and analyzed their single-cell transcriptome using SORT-seq^65^. Autofluorescent cells (viability dye eFluor780-high) and non-EECs were removed to create an EEC-centric dataset (Extended Data Fig. 6a-c). Clustering revealed presence of EEC progenitors (*NEUROG3+)* and mature peptide-hormone producing X, D and L cells (Fig. 6e–f, Extended Data Fig. 6d). We could not resolve ECL and EC cells into separate populations.

**Fig. 6:**
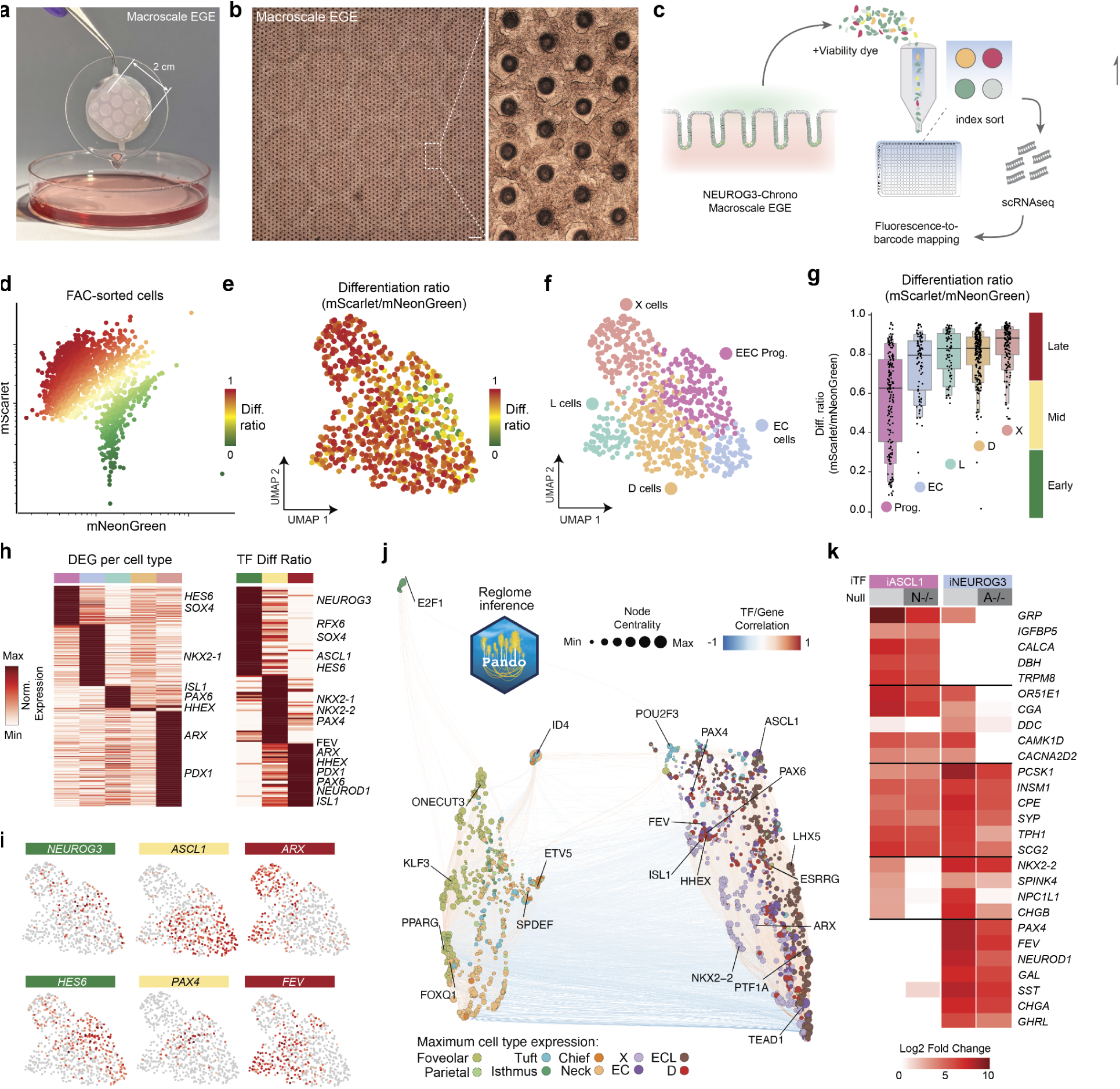
Macroscale EGE resolve EEC differentiation dynamics. **a**, Photograph of a large-scale engineered corpus tissue taken from the 10 cm dish below. **b**, Brightfield images of large-scale EGE at 3 weeks of culture. Scale bar overview: 1 mm, insert: 100 μm. **c**, Workflow of the NEUROG3-Chrono experiment: mEGE were cultured for 2-3 weeks, dissociated and sorted for NEUROG3-Chrono+ cells into 384-well plates. The mNeonGreen and mScarlet fluorescent intensities were registered via index sort. Single cells were sequenced via SORT-seq and fluorescent intensities were mapped back onto the single cell barcodes. **d**, FACs plot showing the expression of mNeonGreen and mScarlet fluorescence intensities of sorted NEUROG3-Chrono+ cells. Each dot represents one cell. **e**, UMAP showing the differentiation ratio (mScarlet/mNeonGreen intensity) of NEUROG3-Chrono+ sorted cells. **f**, UMAP showing the populations of EEC subtypes derived from NEUROG3-Chrono+ sorted cells. **g**, Boxenplots showing the differentiation ratio of EEC subtypes derived from NEUROG3-Chrono+ sorted cells. (**h**) Heatmap showing DEGs between EEC subtypes derived from NEUROG3-Chrono+ sorted cells (left) and TFs that correlate significantly with differentiation ratio, ranked from early (0.0-0.3), mid (0.3-0.6) to late (0.6-1.0) differentiation ratio. (**i**) UMAPs showing expression of selected TFs expressed in EEC progenitors: *NEUROG3*, *ASCL1*, *ARX*, *HES6*, *PAX4* and *FEV*. (**j**) Inferred gene regulatory network using Pando. Transcription factors (TFs) and target genes are represented as nodes scaled by network centrality. Edges between TFs and target genes are colored by the maximum region-to-gene (R2G) importance score. Target genes are colored by the cell type in which they show highest expression; TF nodes are labeled. (**k**) Heatmap showing fold change in gene expression upon ASCL1 overexpression (iASCL1) in wild type and NEUROG3 null (-/-) cultures, and NEUROG3 overexpression (iNEUROG3) in wild type and ASCL1 null (-/-) cultures (n=3-6 EGE per condition).

*NEUROG3*+ progenitors displayed the lowest differentiation ratio, indicating fidelity of the reporter (Fig. 6g). ECs displayed significantly faster differentiation kinetics compared to other EECs (Fig. 6g). To explore TFs acting in the various EEC trajectories, we ranked their expression according to cell type and differentiation ratio (Fig. 6h-i, Supplementary Table 7). Early EECs were enriched for *HES6* and *SOX4*, which have been described in early intestinal EECs^66^. Mid. EECs were enriched for *ASCL1*, *PAX6* and *PAX4*. These TFs were predominantly expressed in EC, L and D cells, respectively. Late EECs expressed *ARX, PROX1, PDX1* (X cells), *HHEX* (D cells) and *ISL1* (L cells). Some of these TFs have been implicated in differentiation of intestinal or pancreatic endocrine cells. For example, *HHEX* regulates expression of somatostatin in pancreatic delta cells^67^, whereas Isl-1-mutant mice lose intestinal GLP-1^67,68^.

Beyond dynamic expression of fate-determining TFs, the interplay with chromatin accessibility determines how gene-regulatory networks shape differentiation and cell states. We thus performed paired scRNA- and ATAC-seq on adult human corpus tissue. We enriched the dataset for rare lineages using unique surface markers CD200 (EECs) and KIT (tuft cells), yielding 951 high-quality EECs and 103 tuft cells (Extended Data Fig. 7a). All mature epithelial lineages were resolved, including chief, foveolar, neck, endocrine, tuft and parietal cells. Additionally, we identified fibroblasts and several immune populations (B, T cells and macrophages). Genes known to be highly expressed in specific gastric lineages showed increased accessibility in the corresponding cell types (Extended Data Fig. 7b-c, Extended Data Fig. 8a-d). This concordance between chromatin accessibility and known marker gene expression confirms the robustness of the ATAC data. Additionally, we observed marked lineage-specific correlation between motif accessibility and expression of PPARG and SREBF2 in foveolar cells and AR in chief cells (Extended Data Fig. 8d). Notably, the orphan nuclear receptor ESRRG, recently linked to parietal cell differentiation^76^, showed unexpectedly broad enrichment across multiple EEC subtypes, pointing to a wider role in gastric endocrine specification.

To identify transcriptional regulators of gastric cell identity, we inferred a gene regulatory network (GRN) from paired single-nucleus RNA- and ATAC-seq data using Pando, linking chromatin accessibility at conserved regulatory elements to target gene expression across epithelial populations^69^ (Fig. 6j, Supplementary Table 8, Extended Data Fig. 9a-b). This analysis revealed both known and previously unrecognized regulators of gastric lineage identity. *POU2F3* activity was restricted to tuft cells, in line with its known role in the intestine and lung - and the tuft cell induction we observed with overexpression of its co-activator (Extended Data Fig. 9c-f, Extended Data Fig. 4f). Various Kruppel-like factors enriched in foveolar cells (Extended Data Fig. 9d-f). *FOXQ1* showed high predicted activity in neck and foveolar cells. In mice, this TF is essential for mucin granule formation in foveolar cells^71^. Neck and chief cells were enriched for *SOX9* and *SPDEF*, which are essential for the differentiation of these lineages in mice^72,73^. EECs and to a lesser extent tuft cells demonstrated activity of *SOX4* (Extended Data Fig. 9f). *SOX4* is sufficient to drive tuft and EEC differentiation in the murine intestine^74^, but has not been linked to gastric secretory cell differentiation. Among EEC subtypes, the GRN resolved subtype-specific regulons: ASCL1 in EC and ECL cells; PTF1A and LHX5 in ECL cells; ARX, ISL1, NEUROD1 and NEUROD2 in X cells; and HHEX and PAX6 in D cells - largely consistent with their expression dynamics in the NEUROG3-Chrono data (Extended Data Fig. 9f, Fig. 6h-i, Supplementary Table 7).

Differentiation of intestinal EECs critically depends on NEUROG3, yet the murine stomach does not lose all EECs upon knockout of *NEUROG3*, maintaining X, EC and ECL cells^77^. Previous work has pointed to ASCL1 as another master TF regulating gastric EEC differentiation in animal models^78–80^, potentially acting upstream of *NEUROG3*^78^. Nevertheless, *ASCL1* expression in our EEC atlas is depleted from NEUROG3+ progenitor cells (Fig. 6c). To resolve the transcriptional targets of these TFs in gastric epithelial cells, we transiently overexpressed *NEUROG3* or *ASCL1* and performed bulk RNAseq. *NEUROG3* overexpression upregulated markers of multiple EEC lineages, including their hormone genes (Fig. 6k, Supplementary Table 9 and 10). Instead, *ASCL1* overexpression was only associated with the induction of EC cells, whereas none of the peptide-producing EEC subtypes were significantly affected. This is consistent with the predicted activity of *ASCL1* in the EC lineage and its exclusion from other EEC subtypes. NEUROG3 overexpression failed to rescue the expression of *DDC*, the enzyme converting 5-hydroxytryptophan into serotonin in ASCL1-null organoids. The expression hierarchy and gain-of-function experiments suggest that *NEUROG3* is a master EEC regulator in the stomach, while *ASCL1* is a key driver of EC identity. Overall, our single cell genomic atlas provides a first comprehensive roadmap into gastric differentiation, including rare EECs, which can serve as the basis for functional exploration of their differentiation.

## Discussion

Here, we have combined bioengineering, genetic engineering and single-cell genomic approaches to build and investigate a homeostatic model of the human gastric epithelium. We assessed selected growth- and transcription factors that regulate cellular diversity. By leveraging the self-organizing and renewal capacities of these epithelial cells, we showcase robust gland-pit organization and longevity in EGE. To our knowledge, there are currently no other gastrointestinal *in vitro* systems beyond EGE that combine a complex cellular composition with unperturbed, long-term growth. Such a model could provide unique insights in processes that require long timescales, such as clonal competition during cancer evolution.

EGE homeostatically generate previously understudied gastric lineages, such as tuft cells and EECs. Importantly, we identified a novel gastric cell type that is *FOXI1*-regulated and transcriptionally resembles ionocytes found in other tissues. These cells express some TFs enriched in tuft cells, such as *POU2F3* and *POU2AF2*, suggesting a shared developmental origin, as has been proposed in the lung. A notable difference with the well-characterized pulmonary ionocytes is the lack of *CFTR* expression enrichment in gastric ionocytes versus other lineages. Instead, *CFTR* is more broadly expressed by neck cells, which secrete both mucus and bicarbonate to neutralize the pH of the mucus layer, forming an alkaline barrier between the epithelium and the acidic lumen. Gastric ionocytes are enriched for V-ATPases and sodium channels, which execute apical ion exchange and various metabolic enzymes required to sustain high energy-demanding processes. These features suggest that gastric ionocytes are equipped to sense and modulate the physical properties of the gastric mucus layer.

Human ASC-derived organoids additionally fail to produce representative numbers of EECs, and even overexpression of *NEUROG3* does not generate the full spectrum of EECs *in vitro*. EGE homeostatically produce all EEC subtypes, including histamine-producing ECL cells. Since EECs are rare and can be long-lived^81^, it has been challenging to capture their transient progenitor states in previous scRNAseq efforts. To address this, we employed NEUROG3-labeling and large-scale EGE cultures to generate an atlas of gastric EECs and their progenitors.

Surprisingly - beyond the well-described EEC lineages - we identified rare L cells *in vivo* and *in vitro.* While these predominantly express the satiety hormone *PYY*, we found a subset expressing the precursor hormone to GLP-1 (*GCG*). These cells additionally produced the protease (PCSK1) required for processing this preprotein into GLP-1, and we identified GLP-1+ cells in tissue. A previous report found glucagon peptides in the human stomach using bulk tissue mass spectrometry, yet the lack of the *PCSK2* expression would render gastric L-cells incapable of significant glucagon production^63^. Given the L-cell rarity compared to the intestinal tract, it is unlikely gastric GLP-1 has similar, systemic functions to distal GLP-1 (e.g., such as the incretin effect). Instead, gastric GLP-1 may signal locally to influence gastric function: previous studies have identified GLP-1 receptors on parietal cells as well as mucinous cells^82–84^, though this GPCR protein has been notoriously difficult to detect with antibody staining. Both our scRNA atlas and spatial transcriptomics data indicate abundant GLP-1 receptor expression among EECs - predominantly X cells - as well as Tuft cells and ionocytes. It may thus be possible that GLP-1 affects local secretion.

The potential applications of EGE are manifold: their scalable nature makes them amenable to large-scale genetic manipulation, for instance in CRISPR screens for regulators of cell type specification or drivers of gastric cancer. The long lifespan allows modeling of clonal dynamics and other emerging properties such as cell type patterning. Furthermore, the apical and basolateral accessibility of the cultures enables cocultures with bacteria or tissue-resident immune cells, and the improved tissue architecture and cell polarization could facilitate drug safety and efficacy studies, such as CLDN18.2-targeting immunotherapies. Finally, we envision mechanical perturbations of the tissues may shed light on the epithelial response to gastric distension. The widespread expression of *PIEZO* channels in gastric EECs suggests inherent mechanosensation - indeed, *PIEZO* channels in murine gastric EECs have been shown to regulate ghrelin secretion and modulate satiety. Together, EGE may support a wide range of research areas in gastric research, from fundamental to therapeutic, and molecular to tissue-level biology.

## Acknowledgments

We thank Justine Fidelin and Kim Schneider from the Roche pRED 360 Labs and Balazs Banfai from the Roche Computational Sciences Center of Excellence for their support in the proteomics and spatial transcriptomics workflow, and Carla Pou Casellas for sharing the integrated ionocyte tissue scRNAseq atlas.

## Funding

H.C. was supported by the European Research Council (ERC Advanced Grant GutHormones 101020405).

## Author contributions

N.A., M.N. and J.B. conceived the study. J.B., N.G. and J.G.C supervised the research. N.A. established corpus organoids. M.N. developed organ-on-chip systems. N.A. and M.N. established culture conditions and performed all chip culture experiments. N.A. and M.N. performed cell dissociation for single cell experiments. N.A. performed flow cytometry, immunofluorescence microscopy, ELISAs and RNA extraction. L.A. preprocessed and analyzed all sequencing modalities including bulk, single-cell and spatial transcriptomics. L.A. integrated scRNAseq atlases, performed benchmarking of chips models, spatial ligand analysis, ionocyte and multimodal integrative analysis. L.G.T. executed library preparation from EGE and tissue. A.C., R.A.S. and J.T.B performed the Xenium workflow. A.V. assisted in Xenium QC and preliminary analysis. Y.B. designed constructs for reporter lines. N.A., J.Bo. and S.R.V. generated reporter lines. J.Bo. performed qPCR. F.V. performed proteomics of culture supernatants, QC and analysis. I.C. generated FFPE blocks and sections of corpus and antrum tissue and performed multiplexed immunofluorescent stainings. N.A., M.N. and L.A. assembled the figures. N.A., J.B., M.N., N.G., L.A. and J.G.C. wrote the manuscript. N.A., M.N., L.A., L.G.T., I.C., A.C., F.V., R.S. and J.T.B. contributed methods. All authors read, edited and approved the final manuscript.

## Competing interests

All authors are employees of Hoffmann-La Roche AG. The company provided support in the form of salaries for authors but did not have any additional role in the study design, data collection and analysis, decision to publish or preparation of the manuscript. We acknowledge additional financial support to H.C. from the European Research Council (Advanced Grant 101020405). M.N. and N.G. are named as inventors on patent applications related to scaffold-guided organoid technology and devices for culturing organoids, filed by The Ecole Polytechnique Fédérale de Lausanne (EP16199677.2, PCT/EP2017/079651, and US20190367872A1) and Hoffmann-LaRoche (EP24167170). J.B. and H.C. are inventors on several patents related to organoid technology. H.C.’s full disclosure is given at https://www.uu.nl/staff/JCClevers/AncillaryActivities.

## Data, code, and materials availability

Plasmids of all reporter constructs, guide RNAs and overexpression vectors will be uploaded on Addgene. All code used for single-cell analysis and data presentation is available via GitHub at https://github.com/devsystemslab/gastric_tissue_lineage_maturation. The accession numbers for the mRNA sequencing data reported in this paper are ArrayExpress E-MTAB-16678 (ECT), E-MTAB-16676 (human developing gastric corpus), E-MTAB-16675 (paired mRNA- and ATAC-seq). Spatial transcriptomics data was deposited at Zenodo (10.5281/zenodo.17953268). Please contact the authors for inquiries. All raw mass spectrometry and Spectronaut search files were deposited in the Proteomics Identifications Database (PRIDE, identifier: PXD074556).

## Methods

### Experimental model and study participant details

Human stomach corpus and antrum tissue samples were obtained within the framework of the non-profit foundation HTCR (Munich, Germany), including informed patient consent. The framework of the HTCR Foundation has been approved by the ethics commission of the Faculty of Medicine in the Ludwig Maximilian University (no. 025-12) and the Bavarian State Medical Association (no. 11142). Four adult corpus (Donor 1: Female, 62 years old, Donor 2: Male, 70 years old, Donor 3: Female, 64 years old, Donor 4: Male, 64 years old), two adult antrum tissues (from Donor 2 and Donor 3, matched) were surgically removed for various oncological indications. Micro- and/or macroscopically tumor-free regions of resections were used for further preparation. One human developing corpus sample (Donor 5: Male, 19 weeks gestation) was obtained after elective pregnancy termination and informed written maternal consent from certified procurement services. The donor consented to the research use of the tissue without restrictions. The tissue was fully anonymized, and all experiments were performed in accordance with relevant guidelines and regulations.

### Organoid establishment and culture

Corpus and antrum organoids were established as described previously^85^. In brief, the basal muscularis serosa and fat were removed using forceps and scissors. The mucosa was minced and dissociated to cell clumps of 5-25 cells using 20 μg/mL Collagenase IV (Sigma Aldrich) and 10 μM dithiothreitol (DTT) in Dulbecco’s Modified Eagle Medium (DMEM, both Thermo Fisher Scientific), strained using 100 um cell strainers (Greiner Bio-One) and plated in reduced-growth factor Matrigel (Corning) domes. Culture medium consisted of Advanced DMEM/F12, supplemented with B27 - vitamin A, Glutamax, HEPES, 100 U/mL Penicillin-Streptomycin (all Thermo Fisher Scientific), 100 mg/mL Primocin (Invivogen) and the following growth factors: 2% Rspo3-conditioned media, 2% Noggin-conditioned media, 0.15 nM WNT-surrogate (all ImmunoPrecise Antibodies), 500 nM A83-01 (Tocris), 50 ng/mL FGF-2 (Thermo Fisher Scientific), 100 ng/mL IGF-1 (BioLegend), 10 nM Gastrin (MedChemExpress), 1 mM N-acetylcysteine (Sigma Aldrich). Media was replaced twice a week, and organoids were passaged by enzymatic dissociation using TrypLE Express (Thermo Fisher Scientific) every week at a 1:10 ratio. After passaging, the medium was supplemented with 10 μM ROCK-inhibitor Y-27632 (Tocris). For details on all medium components used in this study, see Extended Data Table 2.

### Generation of fluorescent reporter lines

Targeting vectors for H2BC4-mTurquoise2, GHRL-P2A-mTagBFP2-PGK-Zeo, LIPF-P2A-NLS-AausFP1-PGK-Puro, ATP4A-P2A-mCherry-PGK-Hygro, GKN1-P2A-NLS-iRFP670-Hygro, NEUROG3-P2A-NLS-mScarlet-P2A-NDegron-NeonGreen for homology-directed repair (HDR) were designed as described previously^23^. Human corpus organoids (Donor 1) were incubated with 1.25% DMSO V/V and 10 μM Y-27632 (Tocris) in culture media 24h prior to electroporation. Organoids were harvested, dissociated using TrypLE Express, 10 μM DTT (both Thermo Fisher Scientific) and 10 μM Y-27632 to clumps of approximately 5 cells. 10×6 cells were resuspended in High Performance Electroporation Solution (BTXpress) containing up to 20 μg DNA at a 1:3 molar ratio of respective pCMV-SpCas9-U6-gRNA expression vector and targeting vector. Cells were transfected using the NEPA21 Super Electroporator (Nepagene) with settings as described before^86^. H2B-mTurquoise2+ organoids could be identified by fluorescence signal and were clonally isolated 14 days post transfection. Organoids successfully modified with cell-type reporters were not identifiable due to lack of differentiation in Matrigel domes. Edited clones were positively selected using 10 μg/mL puromycin (Invivogen) for 7 days, 100 μg/mL hygromycin (Invivogen) for 2 weeks or 100 μg/mL zeocin (Invivogen) for 2 weeks. Surviving clones were screened for site-specific integrations by Sanger sequencing of PCR-amplified genomic DNA. Between 3 and 10 clones with site-specific integrations were pooled to generate polyclonal lines. The antibiotic resistance and iCasp elements were removed by recombination of the LoxP sites flanking these sequences, induced through 24h incubation with 1:100 cell-permeant TAT-CRE Recombinase (Millipore). Non-recombined clones were negatively selected by induction of the iCasp element using 10 μM AP20187 (MedChemExpress). Successful recombination was confirmed through Sanger sequencing of PCR-amplified genomic DNA and 7-day incubation with puromycin, hygromycin or zeocin.

### Generation of NEUROG3 and ASCL1 null mutants

Wild-type organoids were prepared for electroporation as described above. Organoids were transfected with 10 μg pCMV-SpCas9-U6-gRNA and 5 μg gRNA encoding plasmid. Clones were isolated 10 days post transfection and mutant clones were confirmed by Sanger sequencing of PCR-amplified genomic DNA. At least 3 successfully modified clones were pooled to generate polyclonal lines.

### Lentiviral transduction of inducible transcription factor ORF constructs

TRE3G-FOXI1-P2A-3xNLS-mPapaya, TRE3G-POU2AF2-P2A-3xNLS-mScarlet3, TRE3G-NEUROG3-P2A-NLS-mCerulean-P2A-BlasticidinR and TRE3G-ASCL1-P2A-NLS-mCerulean-P2A-BlasticidinR lentiviral constructs were produced by VectorBuilder. Corpus organoids were dissociated to 1 to 5-cell fragments and transduced overnight at 2 TU/cell in organoid medium supplemented with 10 μM ROCK-inhibitor Y-27632 (Tocris) and 5 ug/mL Polybrene (VectorBuilder). Cells were centrifuged at 500xg for 5 mins and plated in reduced-growth factor Matrigel (Corning). After polymerization, domes were covered with organoid culture medium + 10 μM Y-27632. iNEUROG3 and iASCL1 organoids were selected with 5 ug/mL Blasticidin for 2 weeks starting at 3 days post transduction. iFOXI1 and iPOU2AF2 organoids were exposed to 1 μg/mL doxycycline (Sigma) at day 6 post transduction, after which 5-10 pure mPapaya+ or mScarlet3+ clones were picked, pooled and expanded to create polyclonal lines. For overexpression experiments, organoids were cultured in triplicate wells per condition and exposed to 1 μg/mL doxycycline or control media for 7 days. Samples were lysed, processed and sequenced independently.

### Fabrication of biomimetic microtopography hydrogels

For fabrication of hydrogels containing biomimetic microtopography, we used engineering expertise established previously^27^. Stamps featuring corpus gland-mimicking microtopography were fabricated from PDMS via replica molding of 2PP-printed masters. First, arrays of corpus gland-mimicking micropillars were designed in Autodesk Fusion. Several designs were tested and optimized; the final microtopography consists of slightly conical pillars ranging from 70 to 150 µm in diameter (top to bottom) and 750 µm in height. The 3D models were printed using a NanoOne (UpNano) 2PP printer on glass coverslips with UpPhoto resin and a 10×/0.4 NA objective. To generate a master mold with inverted topography, the printed structures were plasma-activated and silanized overnight using vapor-phase trichloro(1H,1H,2H,2H-perfluorooctyl) silane (Sigma-Aldrich). PDMS (Sylgard 184, Dow Corning; 10:1 base-to-curing agent ratio) was poured over the master, degassed under vacuum, and cured at 80 °C for 12 h. The resulting PDMS replica was cut, plasma-activated, and silanized overnight as above. This PDMS negative was then used repeatedly as a master mold for casting final PDMS stamps using the same procedure (Sylgard 184, 10:1 ratio, cured at 80 °C for 12 h).

For housing the hydrogel, we designed a chip containing a central hydrogel chamber flanked by two gel-injection ports and one top reservoir above the hydrogel. The hydrogel precursor was injected through one of the loading ports leading to the central gel chamber. A cylindrical stamp with microtopography on its base was placed on a stamp-holding ledge of the top reservoir before gel injection. The diameter of the open, microtopography-patterned hydrogel area is 4 mm, which, after cell seeding, becomes fully covered edge-to-edge by the epithelium, establishing a tight barrier. After gel polymerization (see details below), the stamp was removed, leaving an inverse microtopography (i.e., an array of microwells) on the hydrogel surface. The top reservoir was then filled with PBS or cell culture medium. The top reservoir was used for cell seeding and subsequently served as the apical medium reservoir.

For scalable chip manufacturing, we developed a cassette-like mold for PDMS injection and curing. Molds were manufactured by CNC milling of acrylic or steel, depending on the intended production scale. Each mold consisted of two parts bearing the inverse topography of the chip’s bottom and top surfaces. When assembled, both parts fit precisely, enabling accurate alignment of both sides of the device. PDMS was injected into the central cavity of the mold, degassed in a vacuum chamber, and cured in an 80 °C oven. After curing, the mold was opened, and the PDMS sheets were cut and O₂ plasma-bonded. After bonding, parts were baked in the oven at 100 °C for at least overnight. Holes in each chip and individual chips were then cut from the large PDMS sheets using punchers. The resulting chips were sterilised in 70% ethanol and air-dried before use.

To generate the extracellular matrix (ECM) in the chip, bovine TeloCol-6 Type I collagen solution (5225-50mL, Advanced Biomatrix) was neutralized following the manufacturer’s instructions and then mixed with Matrigel (356231, Corning) at an 80/20 (v/v) ratio. A PDMS stamp with a crypt-mimicking array of pillar topography was placed in the central top reservoir. After gel injection through the gel-loading port, the hydrogel was polymerized at 37 °C for 15 min. After ECM polymerization, all chip reservoirs were topped up with PBS or cell culture medium. Prepared chips with hydrogels can be stored for multiple weeks at +4 °C or kept in the incubator before use.

Hydrogel stiffness was measured using an MCR 702e MultiDrive oscillatory shear rheometer (Anton Paar) in a parallel plate configuration with a PP25/P3 measuring plate. The Young’s modulus (E) was estimated from the storage modulus (G′), assuming the hydrogel is elastic and nearly incompressible, using the relation E= 2G(1 + ν) ≈ 3G′, where Poisson’s ratio (ν) is approximately 0.5. Across multiple batches and replicates, the hydrogels yielded a Young’s modulus of 4000 ± 500 Pa.

### Seeding and culturing of EGE

Organoids were released from Matrigel 7 days post passaging by incubation in Cell Recovery solution (Corning) for 15 mins on ice. Falcons were centrifuged at 700xg for 5 mins, 4C. Pellets were resuspended in TrypLE + 10 μM DTT + 10 μM Y-27632 + DNase I (Roche) and incubated at 37C for 5 mins. Falcons were vortexed thoroughly, inspected for successful fragmentation under a wide-field microscope and incubated an additional 3 mins if required and triturated with P1000 pipettes to obtain single cells and fragments of max 5 cells. The suspension was strained through 40 μM strainers (Greiner Bio-One) and washed with DMEM + 0.2% BSA (Miltenyi). Cells were centrifuged at 700xg for 5 mins, 4C. Pellets were resuspended in organoid culture medium without A-083 + 10 μM Y-27632 at a concentration of 40 million cells/mL. Chips were dried carefully by aspiration. 4-5 uL of cell suspension was placed at the center of each chip. Chips were incubated without additional medium for 1 hour to allow sedimentation and attachment of the cells to the hydrogel. After incubation, chips were carefully removed from the plastic dishes using a thin spatula and tweezers. 1.5 mL of organoid culture medium + 10 μM Y-27632 was placed in each well of a 12-well plate (CellStar) and chips were placed to float on top of the medium, allowing access of medium from the basolateral side without disturbing the recently seeded cells. After overnight incubation, the chips were submerged in the medium and flipped upside down to facilitate sedimentation of unattached cells. Cells formed confluent epithelia within 2 days, at which point chips were transferred to 6-well plates and placed on an orbital shaker set to 100 RPM, enabling efficient nutrient, oxygen and metabolite diffusion and removal of shed cells. The medium was replaced twice a week.

### Immunostaining and microscopy

EGE were fixed with 4% Paraformaldehyde (PFA) overnight at 4C and washed 3x with DPBS (Dulbecco). EGE were removed from the chips using a 4 mm biopsy puncher (KAI). For imaging of the pit-gland axis, sections of 2-3 rows of glands were cut using a custom mini-guillotine device harboring a razor blade. For immunostaining, EGE sections were permeabilized for 1h with DPBS + 0.5% TritonX + 0.1% BSA, blocked for 2h with DPBS + 0.2% TritonX + 0.5% BSA at room temperature and incubated with primary antibodies overnight at 4C. Sections were washed 3x with DPBS and incubated with secondary antibodies overnight (Extended Data Table 3 for details on antibodies used in this study). After 3 washes with DPBS, sections were transferred to glass-bottom dishes (Ibidi) and overlaid with DeepClear solution (CelExplorer). Corpus and antrum tissues were fixed with 4% PFA overnight 4C. Fixed tissues were embedded in paraffin and sectioned on a Thermo Microm HM355S. Multiplexed immunofluorescence staining of Formalin-fixed paraffin-embedded (FFPE) slides was performed using Ventana Discovery Ultra automated tissue stainer (Roche Tissue Diagnostics, Tucson AZ USA). Imaging was performed on Leica Stellaris 8, using a 25x water-immersion objective (HC FLUOTAR L VISIR 25x/0.95 WATER).

### Secretome preparation and analysis

For all secretome experiments, cultures were washed thoroughly with DPBS 3x before the start of the assay and incubated for 1h at 37C to remove debris and previously secreted proteins. For sampling of the apical secretome, cultures were transferred apical-side up to 12-well plates containing 500 µL DPBS+glucose+calcium (Thermo Fisher Scientific) on the basolateral side and 100 µL on the apical side. Cultures were incubated at 37C for 6h, after which apical medium was collected in 1.5 mL Eppendorfs and processed for LC-MS. For quantification of the hormone secretion capacity, cultures were placed in 500 µL DPBS+glucose+calcium (control) or identical medium supplied with 10 μM Forskolin (Tocris) and 10 μM IBMX (MedChemExpress) for 6h. Basolateral supernatant was collected in 1.5 mL Eppendorfs, centrifuged for 5 mins at 1000xg. Supernatants were aliquoted and stored at −20C until time of analysis. ELISAs for Ghrelin, Histamine (both Abcam), Somatostatin (Novus Bio) and Serotonin (LDN) were carried out according to the manufacturers’ instructions. For pepsin activity quantification, cultures were placed in identical basolateral medium as above (+/-Forskolin). The apical medium consisted of DMEM with pH 7 or adjusted to pH 4 using citrate buffer. Apical supernatant was collected after 6h and was analyzed using a fluorimetric Pepsin/Pepsinogen Assay Kit (Abcam).

### Proteomics of EGE apical secretome

Supernatant samples were precipitated by the addition of ice-cold acetone at a 1:4 v/v ratio. The samples were incubated at −20°C for 48 hours. Precipitated proteins were pelleted by centrifugation at maximum speed for 10 min. The supernatant was discarded, and the protein pellets were briefly air-dried. Pellets were resuspended in 200 µL of PreOmics lysis buffer and heated at 95°C for 10 min. Protein concentration was determined using a BCA protein assay, and sample amounts were normalized to 10 µg of input material. Sodium dodecyl sulfate (SDS) was added to the samples to a final concentration of 5%, followed by a second incubation at 95°C for 10 min. Proteins were reduced, alkylated, and digested using an automated robotic protocol as previously described^87^, with the modification that tryptic digestion was performed offline overnight. Following digestion, the samples were acidified by the addition of formic acid to a final concentration of 2% (v/v). The digests were transferred to a mass spectrometry sample plate and peptides were dried down. Finally, samples were resuspended in resuspension buffer (2% acetonitrile, 0.5% formic acid in water) prior to LC-MS analysis, and a total of 200ng was loaded on column.

Peptide separation was performed using a Vanquish Neo UHPLC system (Thermo Fisher Scientific) coupled to an Orbitrap Astral mass spectrometer (Thermo Fisher Scientific). The system was operated in a trap-and-elute configuration. Peptides were loaded onto a trap column (300 µm i.d. × 5 mm length) at a maximum pressure of 800 bar. Separation was achieved on a 15 cm Aurora Series emitter column with a 75 µm internal diameter packed with C18 resin (IonOpticks, Fitzroy, Australia). The autosampler temperature was maintained at 7°C, and the column was maintained at 50°C. The mobile phases consisted of solvent A (water with 0.1% formic acid) and solvent B (acetonitrile with 0.1% formic acid). Peptides were separated using a 14-minute method with a non-linear gradient. During the separation phase, the flow rate was maintained at 400 nL/min. The gradient profile was as follows: from 2% to 6% solvent B over 0.1 min, 6% to 8% solvent B over 0.8 min, 8% to 30% solvent B over 8.0 min, and 30% to 45% solvent B over 2.0 min. This was followed by a wash where solvent B was increased to 99% over 0.5 min and held for 2.5 min. The instrument was operated in data-independent acquisition (DIA) mode. A full MS scan was acquired in the Orbitrap analyzer over a mass range of *m/z* 380 to 980 at a resolution of 240,000. The normalized automatic gain control (AGC) target was set to 500%, and the maximum injection time was 3 ms. DIA scans were acquired in the Astral analyzer over a mass range of *m/z* 380 to 980 using 150 isolation windows of 4 *m/z* width with no overlap. The normalized collision energy (HCD) was set to 25%. The normalized AGC target for DIA scans was 200%, and the maximum injection time was set to 5 ms. Raw data were processed using Spectronaut (v. 20.2.250922.92449), without a spectral library. Data was searched against the human Uniprot database (as of January 2022). Default settings for the directDIA workflow were applied with 1% FDR at PSM, Peptide and Protein Group levels, as well as the following deviations: decoy generation method was set to ‘Inverse’, Protein LFQ Method was set to ‘Quant 2.0 (SN Standard)’, and Major and Minor Group Quantity were set to ‘Median’. Downstream data analysis was performed in R (v 4.4.1) using the MSstats package^88^.

### Processing of EGE for single cell applications

EGE were removed from the chips using a 4 mm biopsy puncher (KAI), and incubated with Liberase TH for 10 mins at 37C to dissolve the hydrogel. Macroscale cultures were removed from the hydrogels 2-4 weeks after seeding by scraping the apical surface with forceps, aided by a stereomicroscope. The hydrogel was washed with DPBS + 0.1% BSA until all tissue fragments were detached. The fragments were collected in 50 mL Falcons and centrifuged at 500xg for 5 mins at 4C. The epithelia were dissociated to single cells using Neural Tissue Dissociation kit P (Miltenyi) according to the manufacturer’s instructions. Dissociation reactions were supplemented with 10 μM DTT + 10 μM Y-27632. For FACs and flow cytometry, cell suspensions were incubated with PE/Cyanine7 anti-human CD117 (c-kit) and/or APC anti-human CD200 (both BioLegend) and eFluor 780 Fixable Viability Dye (Thermo Fisher Scientific) on ice for 15 mins. The cells were washed with 10 mL of DPBS, centrifuged at 700xg for 5 mins at 4C. The pellets were resuspended in FACs buffer (DPBS + 0.1% BSA + 0.02 M EDTA (Thermo Fisher Scientific) + 10 μM DTT + 10 μM Y-27632). Samples were analyzed on a Cytoflex S 86 (Beckman Coulter) or sorted on a Aria Fusion (BD Biosciences) or Aurora CS (Cytek). For NEUROG3-Chrono SORT-seq, single cell suspensions were incubated with eFluor 780 Fixable Viability Dye (Thermo Fisher Scientific) on ice for 15 mins. The cells were washed with 25 mL of DPBS, centrifuged at 700xg for 5 mins at 4C. The pellets were resuspended in FACs buffer (DPBS + 0.1% BSA + 0.02 M EDTA (Thermo Fisher Scientific) + 10 μM DTT + 10 μM Y-27632). Single cells were gated on live cells and sorted into 384-well plates on an Aria Fusion (BD Biosciences) and immediately snap-frozen prior to single cell RNA sequencing. Library preparation and single-cell RNA sequencing was performed by Single Cell Discoveries (Utrecht, the Netherlands) according to previously published SORT-seq protocols^65^.

### Processing of tissue for single cell applications

For single cell RNA sequencing, the mucosa tissue from one developing human corpus (Donor 5) was dissected using forceps and scalpels. For multiome sequencing, adult corpus tissue (Donor 4) was processed in a similar fashion. Samples were dissociated to single cells using Neural Tissue Dissociation kit P (Miltenyi) according to the manufacturer’s instructions. Dissociation reactions were supplemented with 10 μM DTT + 10 μM Y-27632. The cells were incubated with anti-CD326 (Ep-CAM), anti-human CD117 (c-kit) (both Biolegend), anti-human CD200 and eFluor 780 Viability dye (eBioscience) for 15 mins at 4C, washed with DPBS (Dulbecco) + 0.1% BSA and pelleted at 500xg, 4C for 5 mins. The pellets were resuspended in FACs buffer (DPBS + 0.1% BSA + 0.02 M EDTA (Thermo Fisher Scientific) + 10 μM DTT + 10 μM Y-27632) and EpCAM+ cells were sorted on a FACS Aria Fusion, enriching for CD200+ and c-kit+ cells.

### Single cell RNA sequencing

Single-cell RNA-seq libraries were generated using 10x Genomics Chromium Next GEM Single Cell 3ʹ Kit v3.1 according to user manual CG000315 Rev E. Briefly, cell concentration, viability and percentage of aggregation of single-cell suspensions were quantified on the automatic cell counter Countess 3 FL (AMQAF2000, Thermo Fisher Scientific) with Trypan Blue Stain (T10282, Thermo Fisher Scientific). For each condition, cells were loaded into one lane of a Chromium Chip G (1000120, 10x Genomics) with a target recovery of 1000 - 10000 cells. Single-cell encapsulation, barcoding and library preparation were performed according to the manufacturer’s guidelines. Libraries were quantified with Qubit 1X dsDNA HS (Q33231, Thermo Fisher Scientific) and Bioanalyzer High Sensitivity DNA Kit (5067-4626, Agilent Technologies) and were sequenced on the Illumina NovaSeq 6000 and NovaSeq X platforms.

### Single-cell multiome sequencing

Nuclei isolation was performed based on 10x Genomics Demonstrated protocol CG000365 Rev C. Single cells were centrifuged for 5 min at 600g and 4°C. The cell pellet was resuspended in cold lysis buffer (10 mM Tris-HCl pH7.4, 10 mM NaCl, 3 mM MgCl_2_, 1% BSA, 1 mM DTT, 0.1% Tween-20, 0.1% Igepal CA-630, 0.01% Digitonin, 1U/µl RNAse inhibitor). After 8 min on ice, cold wash buffer was added (10 mM Tris-HCl pH7.4, 10 mM NaCl, 3 mM MgCl_2_, 1% BSA, 1 mM DTT, 0.1% Tween-20, 1U/µl RNAse inhibitor) and nuclei were spun for 5 min at 600g and 4°C. The pellet was resuspended in cold wash buffer and nuclei were counted with Propidium Iodide (ViaStain AO/PI, CS2-0106-5ML, Nexcelom Bioscience) using the automatic cell counter Countess 3 FL (AMQAF2000, Thermo Fisher Scientific). After spinning, nuclei were resuspended in Diluted Nuclei Buffer (10x Genomics) and their quality was evaluated by microscopy with DAPI and bright-field. Single-cell multiome was performed using 10x Genomics Chromium Next GEM Single Cell Multiome ATAC + Gene Expression reagents and corresponding user guide CG000338 Rev F. One lane of Chromium Next GEM Chip J (1000234, 10x Genomics) was loaded targeting 10.000 nuclei. snRNA-seq and snATAC-seq libraries were sequenced on the Illumina NovaSeq X Plus platform.

### Processing and integration of EGE single cell transcriptomic data

EGE-derived transcript count matrices were aligned to hg38 using CellRanger (v9.0.0, v8.0.1 for Corpus4_OCflip_6wk). Quality control retained cells with ≥100 transcripts, ≥500 genes, ≤60,000 counts, and <50% mitochondrial reads. Mitochondrial and ribosomal genes were removed, and count matrices were normalized and log-transformed. We identified 3,000 highly variable genes using the Seurat v3 method with batch as additional argument. To minimize confounding by mitochondrial content, we quantified gene–mitochondrial percentage correlations and excluded genes with |r| ≥ 0.1 from the initial set of highly variable genes. Data were integrated using scVI (scvi-tools v1.4.0) and clustered with Leiden algorithm (resolution = 2.0). Rare epithelial populations were resolved by subclustering clusters 24, 32, and 33 at resolution 1.0; an aberrant cluster co-expressing enteroendocrine and non-EEC markers was excluded. Cell types were assigned using snapseed with hierarchical marker annotation and manual refinement. For integration and initial cell type calling, both the baseline (called ‘base’ or ‘expansion medium, EM’) and ERBB-ligand-treated culture conditions were used. All combined single cell RNA datasets are deposited. For all visualization and downstream analysis in this manuscript, the EGE dataset was subset to baseline culture conditions only.

### Construction of human stomach reference atlases

We constructed reference atlases for adult and developing human stomach tissue. For the adult reference, we integrated two published datasets ^31^ and ^32^, retaining histologically normal epithelial cells from the first source and including the second in full. Quality control filters required ≥500 transcripts and genes per cell, ≤50,000 total counts, and <20% mitochondrial reads. Data were integrated using scVI ^89^ (scvi-tools v1.4.0), clustered with the Leiden algorithm (resolution = 2.0), and annotated using snapseed followed by manual refinement; a tumor-like population was excluded. For the developing stomach reference, we performed scRNA-seq on human fetal stomach tissue (Donor 5), processed with CellRanger (v6.0.2), and applied quality control criteria (≥500 transcripts, ≥1,000 genes per cell, ≤50,000 counts, <30% mitochondrial reads). Data were normalized, log-transformed, and highly variable genes identified using the Seurat v3 method. We regressed out technical factors (total transcript counts and mitochondrial percentage), performed scaling, PCA and UMAP embedding, and clustered with the Leiden algorithm (resolution = 2.0). Cell type annotation was performed using snapseed with curated marker hierarchies.

### Construction of an integrated human stomach reference

To generate a unified atlas of developing and adult stomach epithelium, we removed all non-epithelial cells from the individual datasets. The two datasets were then merged, and only genes shared across both were retained. To ensure balanced feature selection across developmental stages, we reselected highly variable genes jointly across the combined dataset using the Seurat v3 method (5,000 genes) with batch-aware selection. Integration was performed using scVI^89^ (scvi-tools v1.4.0) and a two layer network (layer normalization on encoder and decoder, covariate encoding enabled, dropout rate = 0.2). After training the scVI model, we refined the embedding using scANVI^90^ to incorporate cell-type supervision. The model was trained for an additional 20 epochs using balanced sampling across labeled categories to obtain a harmonized latent space suitable for downstream clustering and annotation.

### Benchmarking of EGE against adult reference

To benchmark EGE against the adult stomach reference, we restricted both datasets to shared epithelial cell types (ECL cells, Intermediate mucous cells, Foveolar cells, EC cells, X cells, Chief cells, Tuft cells, Isthmus cells, Neck mucous cells, and Ionocytes), merged at the raw count level, and retained only shared genes. We selected 5,000 highly variable genes using batch-aware Seurat v3 selection and integrated the datasets using scVI (scvi-tools v1.4.0) with batch correction. The embedding was refined using scANVI with cell-type supervision initialized from the trained scVI model. Cell type labels were transferred from reference to EGE using CellMapper (https://doi.org/10.5281/zenodo.16606250, v0.2.3), a k-NN-based mapping tool applied to the scANVI latent space. Using these transferred reference annotations, we estimated the percentage of correctly assigned cells per cell type as a proxy for the confidence in core cell type identity. For tuft and ionocyte classification, we fine-tuned the scANVI model on the EGE tuft/ionocyte cluster (100 epochs) and transferred labels using a k-nearest neighbor graph (k = 15), with results summarized as a normalized confusion matrix.

### Comparison with published model systems

We reanalyzed raw count matrices from Aguilar et al.^11^ following standard quality control (≥500 transcripts and genes per cell, ≤50,000 counts, <30% mitochondrial reads). After normalization and log transformation, we identified 2,000 highly variable genes using Seurat v3, scaled the data, and performed PCA. Neighborhood graphs were constructed using 15 nearest neighbors and 40 principal components, followed by Leiden clustering (resolution = 0.5) and UMAP visualization. Epithelial heterogeneity was resolved by subclustering organoid (cluster “1”) and monolayer (cluster “0”) populations at resolution 0.5, with manual annotation based on marker genes. To assess model fidelity, datasets were restricted to shared cell types (Isthmus, Neck, Foveolar, Tuft) and the EGE dataset to 6-week-old standard culture samples. Cosine similarity was computed between mean expression profiles of the top 3,000 highly variable genes from the adult reference and each stomach model, with mean similarity calculated across standard culture conditions.

### Identification of ionocyte-enriched genes

Ionocytes and tuft cells were identified independently in each stomach dataset (in vitro, adult in vivo reference, developing in vivo reference) by subclustering the epithelial compartment and annotating clusters based on established marker gene expression. For each dataset, cells annotated as “Ionocytes” or “Tuft cells” were retained, and analyses were restricted to genes shared across all datasets.

Differential expression between ionocytes and tuft cells was performed separately for each dataset using Scanpy (v1.11.2) (rank_genes_groups) with the t-test and overestimated variance (t-test_overestim_var) on the log-normalized expression matrices. Per-dataset differential expression results were extracted, annotated with dataset identity, and combined into a single summary table for cross-condition comparisons.

In the adult in vivo dataset, we defined *adult ionocyte-enriched differentially expressed genes* as genes upregulated in adult ionocytes compared to adult tuft cells, applying thresholds of p-value < 0.1 and log fold change > 1. To further ensure specificity beyond the closely related tuft cell population, we additionally required that these genes exhibit a positive log fold change when comparing adult ionocytes to all other epithelial cells. Genes meeting both criteria constituted the set of adult ionocyte-enriched DEGs (n = 1,022).

These genes were subsequently annotated based on overlap with previously reported ionocyte-specific differentially expressed genes identified in non-gastric tissues. To define a highly specific stomach adult ionocyte signature, we additionally retained only genes showing highest expression in ionocytes across all three datasets (in vitro, adult in vivo, developing in vivo) and ranked them by their mean tissue-specificity (τ) score across datasets^91^.

### NEUROG3-Chrono analysis

Transcript count matrices were obtained directly from Single Cell Discoveries using their proprietary processing pipeline. The resulting Poisson-corrected count matrices were subsequently imported and merged into a single AnnData object for downstream analysis. Analyses were restricted to protein-coding genes, and gene symbols were made unique. Quality control metrics were computed using Scanpy (v1.11.2). Cells were retained if they contained ≥500 and ≤100,000 total counts and ≥200 detected genes. Genes expressed in fewer than three cells were removed. Cells with >50% mitochondrial transcript content were excluded. ERCC spike-ins, mitochondrial genes (MT-), and ribosomal genes (RPS*, RPL*) were removed prior to downstream analyses.

Compensated .fcs files from the corresponding FACS index sort were processed in R using the flowCore package (v2.16.0). Fluorescence measurements were then linked to the single-cell transcriptome measurements using plate and well identifiers. Alexa488 (Green) and PE-CF594 (Red) fluorescence channels were robustly normalized using 1st–99th percentile min–max scaling and a refined fluorescence ratio was calculated as:

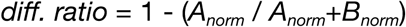

This fluorescence-derived metric was used as a proxy for differentiation time (DT). Autofluorescent cells were identified by clustering in the two-dimensional fluorescence space (diff. ratio vs. log-transformed BV711-A, corresponding to the eFluor 780 viability dye). After scaling, a k-nearest neighbor graph (n_neighbors = 10) was constructed and Leiden clustering was performed (resolution = 0.1).

Library-size normalization (target sum = 10,000) and log-transformation were applied and stored as a dedicated layer. Highly variable genes (n = 2,000) were selected for downstream analyses. Technical covariates (total counts and mitochondrial percentage) were regressed out, and the data were scaled (maximum value = 10). Principal component analysis (50 components) was performed, followed by batch-aware neighbor graph construction using BBKNN^92^ with sample as batch key. UMAP embedding was computed for visualization, and Leiden clustering was performed at multiple resolutions (0.3–2.0). Clusters were then annotated by cell type. Clusters corresponding to high autofluorescence were annotated accordingly. Downstream analysis was restricted to autofluorescence-low EECs. Differential expression across annotated EEC subtypes was performed using Scanpy’s sc.tl.rank_genes_groups function with the *t-test_overestim_var* method, which overestimates the variance within each group to provide more conservative statistics. Genes with fewer than five total counts across all cells were removed prior to testing. Differentially expressed genes were defined as those with log fold-change > 0.5 and adjusted *P* < 0.1 and were exported for downstream analyses.

To identify transcription factors (TFs) that vary along differentiation, cells were stratified according to the fluorescence-derived differentiation metric (*diff_ratio*) using the decoupler package (v2.0.1). Cells were grouped into three ordered bins (Low, Medium, High) based on predefined thresholds (≤0.2, 0.2–0.6, >0.6). Differences in TF activity across these differentiation levels were assessed using one-way ANOVA for each TF. Only TFs with sufficient observations in all three groups were included. TFs with *P* < 0.05 were considered significantly variable and ranked by F-statistic for downstream analyses.

### Bulk RNA extraction, qPCR and RNA sequencing

Hydrogels with corpus epithelia were removed from the chips using a 4 mm biopsy puncher (KAI) and incubated in 350 uL RLT buffer (QIAGEN) for 2 minutes at RT to lyse the cells. The lysates were stored at −20C or processed immediately to RNA using an RNeasy Mini Kit (QIAGEN) with genomic DNA digestion.

For qPCR, cDNA was produced using the Transcriptor First Strand cDNA Synthesis Kit and qPCRs were run on the LightCycler Pro using the SYBR-Green I Master Mix (all Roche). The following primers were used for qPCR analysis: ACTB (Fw: CATGTACGTTGCTATCCAGGC, Rv: CTCCTTAATGTCACGCACGAT), GAPDH (Fw: GGAGCGAGATCCCTCCAAAAT, Rv: GGCTGTTGTCATACTTCTCATGG), FOXI1 (Fw: CGGCCACCCTATTCCTACTC, Rv: TCGGCCACGTACTGGTAGA), ASCL3 (Fw: GGACAACAGAGGCAACTCTAGT, Rv: AGGGGCAGGGTTCACTGTAAT), ATP6V1B1 (Fw: GGCGGTCACCCGAAACTAC, Rv: GGACGATCTCCGCATACTGG), ATP6V0A4 (Fw: CTCCCACGGGAAATGATTACC, Rv: CGTCTCAAAGAAGTCTTGGGTT), SEMA3C (Fw: TTTGCGTGTTGGTTGGAGTAT, Rv: TCCTGTAGTCTAAAGGATGGTGG), TFCP2L1 (Fw: CAGCCCGAGCACTACAACC, Rv: CTCCCAGCTTCCGATTCTCC), POU2AF2 (Fw: GTCACGTCAGGTTACTACGGT, Rv: CCACGCTGGAGGTGTAGAC), TRPM5 (Fw: TTGCTGCCCTAGTGAACCAG, Rv: GCACGATGTCCTCCCAAGAG), BMX (Fw: ACCTGAGGAGTCACGGAAAAGG, Rv: CTGTCCACCAAGCAGTTACGAG), ALOX5 (Fw: ACAAGCCCTTCTACAACGACT, Rv: AGCTGGATCTCGCCCAGTT), AVIL (Fw: ACAACGACCCTGGGATCATTG, Rv: GTCGAGAGGATGACGTAGCAG), POU2F3 (Fw: GCTGGAGAAGTTTGCCAAGACC, Rv: GTGAGATGGTGGTCTGGCTGAA), ATP4A (Fw: CTCACCACCGACGACAATCTG, Rv: TTCCTGGTAGTAGCCAAAGCA), ATP4B (Fw: TGGGTGTGGATCAGCCTGTA, Rv: CTGGTCTTGGTAGTCCGGTG), GKN1 (Fw: CTGTCCACTGCTTTCGTGAAG, Rv: GTCCCATCCGTTGTTATTGTCAA), LIPF (Fw: CTATGCTCTAGCTCCTGTTGC, Rv: CCAAGCGACTCGTGTTAAAGTTC), MUC5AC (Fw: CAGCACAACCCCTGTTTCAAA, Rv: GCGCACAGAGGATGACAGT), MUC6 (Fw: CTGCCCTATACCAGCAATGGA, Rv: CTGACCCATGTACTTCCGCTC). Each sample was analysed as a technical triplicate, Cq values were averaged. Relative gene expression was calculated using the ΔΔCq method.

Library preparation, QC and Illumina sequencing was performed by Alithea Genomics as per the BRB sequencing protocol (BRB-seq Library Preparation Kit for 96, 384, and 1536 Samples).

### Bulk transcriptomics analysis

Alithea Genomics performed read alignment to read alignment and transcript quantification according to the BRB sequencing protocol (https://alitheagenomics.com/services/brb-seq, see <u>BRB-seq Library Preparation Kit for 96, 384, and 1536 Samples</u> for all details). FASTQ files from the same library were merged into single R1 and R2 files prior to analysis. Read quality was assessed using FastQC. Reads were aligned to the human reference genome (GRCh38) using STAR^93^. For multiplexed RNA-seq data, alignments and quantification were performed using STARsolo. Following alignment, per-sample read count matrices and summary statistics were generated from the STARsolo output BAM files using FastReadCounter. Bulk RNA-seq count data and associated metadata were processed by removing lowly expressed genes (total count < 10 across samples). For the iNEUROG3 and iASCL1 induction experiments, Ensembl gene IDs were mapped to HGNC gene symbols using *mygene*, and unmapped or duplicated entries were excluded.

Differential expression analysis was performed using PyDESeq2. After model fitting, pairwise contrasts were computed for each treatment relative to the corresponding control (EM control for the bulk media screen; control for iNEUROG3/iASCL1 induction), and differential expression statistics were obtained using standard DESeq2 inference procedures. Genes with an adjusted *p*-value < 0.05 and |log₂ fold change| > 1 were considered significantly differentially expressed. For visualization, counts were normalized to 1e6 per sample and log-transformed.

### Preprocessing of paired single-cell multiome data

Sequencing data were processed using Cellranger ARC (v2.0.2; 10x Genomics) with default parameters. Reads were aligned to the human reference genome and transcriptome (hg38; 10x Genomics reference, v2.0.2). The resulting fragment and gene count files were used for downstream analyses in SnapATAC2 (v2.8.0) and Scanpy (v1.11.2). Fragment files were imported into SnapATAC2 using a curated hg38 genome annotation, and nonstandard chromosomes were excluded from analysis. Cells with fewer than 1,000 fragments or a transcription start site enrichment (TSSe) score below 7 were removed. For each sample, a 5-kb bin tile matrix was constructed, and 50,000 informative features were selected. Doublets were identified using Scrublet and excluded. Dimensionality reduction was performed using spectral embedding followed by UMAP. Processed ATAC AnnData objects were saved for downstream integration. Single-cell RNA-sequencing (scRNA-seq) data was imported into Scanpy and processed separately. Cells with fewer than 100 transcripts, fewer than 1,000 detected genes, more than 20,000 total counts, or greater than 40% mitochondrial reads were removed, and genes expressed in fewer than three cells were excluded. Count matrices were normalized to 10,000 counts per cell and log transformed. Highly variable genes (3,000 per sample) were identified using the Seurat v3 method with batch correction. The data were scaled to unit variance (values capped at 10), followed by principal component analysis (PCA), construction of a 15-nearest-neighbor graph using 30 principal components, UMAP embedding, and Leiden clustering across multiple resolutions. Parietal and chief cells, together with a small fibroblast cluster, were identified by subclustering Leiden cluster 5. Cell identities were assigned based on canonical marker expression. For downstream integrative analyses, both AnnData objects were subsequently subsetted to retain the same set of matched cells. Gene activity matrices were generated from ATAC fragment files using the make_gene_matrix function implemented in SnapATAC2 and a custom hg38 genome annotation. Genes detected in fewer than five cells were excluded. The resulting matrices were normalized, log-transformed, and smoothed using MAGIC to reduce sparsity and enhance signal structure. For peak-level analyses, chromatin accessibility peaks were called using MACS3 on fragments aggregated by cell type within each sample. Identified peaks were subsequently merged across samples to generate a unified peak set. A peak-by-cell count matrix was constructed using this consensus peak catalog and used for downstream dimensionality reduction and integration analyses. For genome coverage visualization, fragment files and count matrices from both modalities were transferred from Scanpy to Seurat and visualized using the CoveragePlot function implemented in Signac.

### Custom Motif database

Position-weight matrices (PWMs) were obtained from the ArchR (v1.0.3)^94^ motif database and complemented with curated motifs from Pando (v1.1.1)^69^. Motifs from both sources were merged into a unified set, and transcription factor (TF) names were standardized to approved gene symbols. Only motifs with valid and unambiguous gene symbol mappings were retained. In cases where the same TF was represented in both databases, the ArchR-derived motif was prioritized, and the corresponding Pando motif was removed. The resulting non-redundant motif collection was used for all downstream analyses.

### Motif enrichment using chromVAR

To quantify TF activity, motif deviation (chromVAR) scores^95^ were computed using the pychromVAR (v0.0.4) workflow. GC bias and genomic sequence information were incorporated into the peak set, and background peak distributions were estimated to control for technical biases. The curated PWMs described above were matched to accessible chromatin regions using a significance threshold of *P* < 5 × 10⁻⁵. ChromVAR deviation *z*-scores were then calculated for each motif across all cells, providing cell-level TF accessibility activity scores for downstream analyses.

### Gene Regulatory Network inference using Pando

Gene regulatory networks (GRNs) were inferred from stomach multiome (RNA + ATAC) data using Pando (v1.1.1). A Signac object^96^ (v1.16.0), containing gene expression and chromatin accessibility assays generated via SnapATAC2, served as input. The analysis was restricted to protein-coding genes and selected epithelial and enteroendocrine cell populations. RNA counts were log-normalized, and 5,000 highly variable genes were identified. ATAC peaks were restricted to standard hg38 chromosomes, overlapping regions were removed, and a refined chromatin accessibility assay was reconstructed. From this a GRN object was initialized using Pando’s initiate_grn function with the ATAC and RNA assays as input. Exonic regions were excluded, and candidate regulatory regions were restricted to conserved elements defined by *phastConsElements20Mammals.UCSC.hg38* as provided in Pando. Transcription factor binding motifs were then identified in candidate regions using the find_motifs function with the aforementioned curated PFM set and the hg38 reference genome (*<u>10.18129/B9.bioc.BSgenome.Hsapiens.UCSC.hg38</u>*).

GRN inference was performed using GREAT-based peak-to-gene assignment combined with generalized linear models (GLMs), modeling gene expression as a function of TF binding at linked regulatory regions (Pando’s infer_grn function, otherwise default parameters). Models were filtered based on goodness-of-fit and minimum variable contribution criteria (p_thresh = 0.1; nvar_thresh = 2; min_genes_per_module = 1; rsq_thresh = 0.01).

TF modules (regulons) were defined by aggregating significant TF–target interactions and separating positively and negatively associated targets. Regulon activity scores were calculated per cell using Seurat (v5.3.0) AddModuleScore. The resulting TF–target networks were visualized using graph-based approaches, and selected TF-centered subnetworks were extracted for detailed analysis and interpretation.

### *In situ* gene expression analysis sample preparation

#### Custom gene panel design

A custom 480-gene panel was designed for the Xenium *in situ* gene expression analysis platform using the 10x Genomics Cloud Analysis custom panel design tool. Target selection was informed by expression analysis of previously published single-cell RNA sequencing (scRNA-seq) datasets of the human stomach^31^. The panel included cell-type-specific markers for gastric epithelium (parietal, chief, enteroendocrine, tuft, ionocyte and mucous cells), stromal and immune cells, as well as all growth factors and associated receptors expressed in stomach tissue. To avoid optical crowding, targets with high expression (> 50 transcripts per cell) were excluded.

#### Tissue preparation and histology

Human stomach tissue blocks (fundus and corpus regions) were processed following standard clinical pathology protocols. FFPE blocks were sectioned at a thickness of 5um using a Epredia HM355S microtome with cool-cut and section transfer system. One section of the antrum and one section of the corpus were mounted side-by-side on each Xenium Slide within the capture area. Sections were baked at 42°C for 3 hours to ensure tissue adhesion and stored in a desiccator at room temperature prior to library probe hybridization.

#### Xenium in situ workflow

The Xenium In Situ workflow was performed according to the *Xenium In Situ Gene Expression - FFPE Protocol* (10X Genomics CG000749 and CG000582). Briefly, sections underwent deparaffinization using xylene, followed by ethanol rehydration and decrosslinking (target retrieval). The custom probe set was hybridized to the tissue sections overnight, followed by stringent washes to remove unhybridized probes. Ligation was performed to join the two-part probes at the target site, and circularized probes were amplified via rolling circle amplification (RCA).

#### Tissue staining, imaging and cell segmentation

To facilitate cell boundary detection, an optional multi-modal cell segmentation staining was performed using the Xenium Cell Segmentation Add-on Kit. Sections were stained using a cocktail of fluorescently labeled antibodies/rRNA probes (including DAPI for nuclei, ATP1A1, E-Cadherin, and CD45 for cell membrane boundaries, 18S Ribosomal RNA for cytoplasm and Vimentin and Apha-SMA for interior protein) according to the manufacturer’s instructions. Slides were loaded into the Xenium instrument for automated cycles of fluorescent probe hybridization, imaging, and chemical stripping. Data were processed using the Xenium onboard analysis pipeline, which performs image registration, transcript spot calling, and cell segmentation based on the multi-modal staining markers.

#### Spatial transcriptomics analysis and quantification of ligand patterning

Spatial transcriptomics data were imported into the SpatialData (v0.5.0) framework using the Xenium converter. Per-cell statistics from all spatial blocks were merged and jointly analyzed altogether in Scanpy (v1.11.4). Cells with fewer than 10 detected transcripts and genes expressed in fewer than five cells were removed. Counts were normalized by total counts and log transformed. Principal component analysis (PCA) was performed, followed by construction of a nearest-neighbor graph, UMAP embedding, and clustering (resolution of 1.0). Clusters were annotated based on the expression of manually curated marker genes.

To assess ligand expression specifically within the gastric epithelium and quantify it along the epithelial axis, we manually curated epithelial masks for each Xenium block. The main epithelial axis was extracted by skeletonization, and perpendicular cross-sections were constructed along this axis. Each epithelial cell was then projected onto its nearest cross-section to assign a continuous coordinate describing its position along the epithelium.

For each block and each gene in the panel, we then tested for non-linear expression changes along the epithelial axis using generalized additive models (GAMs) (pygam, v0.9.1). A spline model including a smooth function of *axis_position* was compared to a null (intercept-only) model using an F-test. From this we extracted p-values, AIC values, and the predicted expression range along the axis as a measure of effect size. P-values were adjusted for multiple testing within each block using Benjamini-Hochberg correction^97^.

We retained all annotated ligands with a significant adjusted pvalue < 0.05 and a predicted expression range > 0.03 resulting in 25 remaining ligands. Within each block, epithelial cells were ordered by axis position and grouped into 20 bins. Mean expression per ligand and bin was computed to obtain ligand × bin matrices and min max normalized along the bins. For region-level summaries, block-specific profiles were interpolated onto a shared axis grid and subsequently averaged within regions.

## Quantification and statistical analysis

The sample size was not predetermined using statistical methods, and experiments were not randomized, nor were the investigators blinded during experiments and analyses. All data are presented as mean ± standard error of the mean (SEM), unless stated otherwise. Information can be found in the figure legends. Statistical tests included unpaired two-tailed T-test for Figures … using the R package stat_compare_means in RStudio version 4.4.1. ns p > 0.05, ∗ p ≤ 0.05, ∗∗ p ≤ 0.01, ∗∗∗ p ≤ 0.001, ∗∗∗∗ p ≤ 0.0001. Parameters for integration, clustering, differential expression analysis and similarity scoring of bulk, single cell RNAseq and spatial transcriptomics data can be found in the respective methods sections.

---

**Extended Data Fig. 1:**
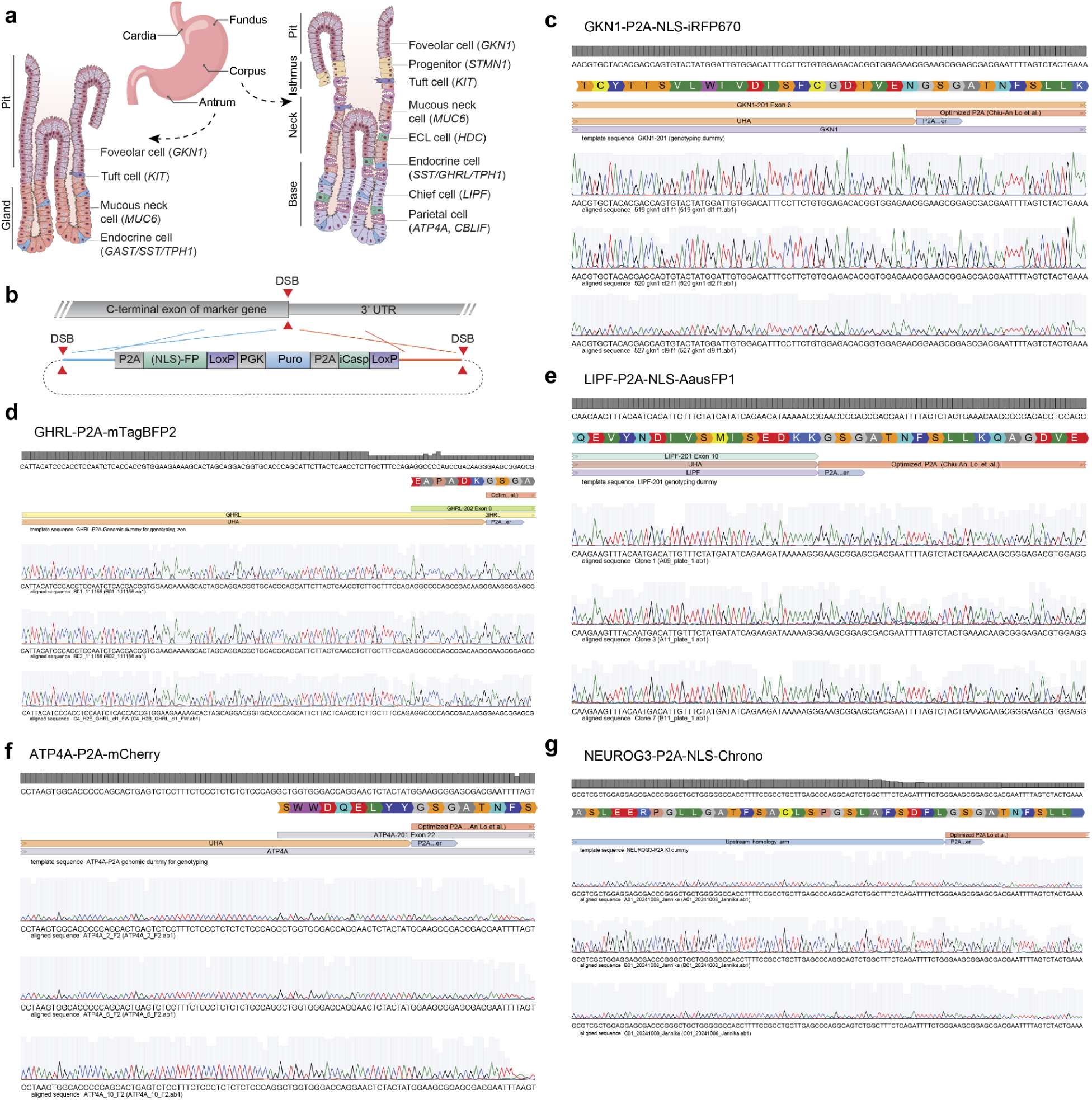
Genetically engineered reporters of gastric epithelial lineages. **a**, Schematic overview of the human stomach epithelium depicting regional and structural organization along the pit–gland axis, including major cell types and representative markers. **b**, Schematic of the fluorescence reporter construct, comprising a P2A-linked (NLS)-fluorescent protein cassette followed by a PGK-driven puromycin resistance gene, a P2A and inducible Caspase module, flanked by LoxP sites enabling cassette removal. The construct is targeted to replace the endogenous stop codon of the target gene. **c**, DNA template of the GKN1 C-terminal exon and P2A preceding the fluorescent protein, showing Sanger sequencing traces of three successfully edited organoid clones below. NLS: Nuclear localization signal. UHA: Upstream Homology Arm. **d**, DNA template of the GHRL C-terminal exon and P2A preceding the fluorescent protein, showing Sanger sequencing traces of three successfully edited organoid clones below. UHA: Upstream Homology Arm. **e**, DNA template of the LIPF C-terminal exon and P2A preceding the fluorescent protein, showing Sanger sequencing traces of three successfully edited organoid clones below. NLS: Nuclear localization signal. UHA: Upstream Homology Arm. **f**, DNA template of the ATP4A C-terminal exon and P2A preceding the fluorescent protein, showing Sanger sequencing traces of three successfully edited organoid clones below. UHA: Upstream Homology Arm. **g**, DNA template of the NEUROG3 C-terminal exon and P2A preceding the fluorescent proteins, showing Sanger sequencing traces of three successfully edited organoid clones below.

**Extended Data Fig. 2:**
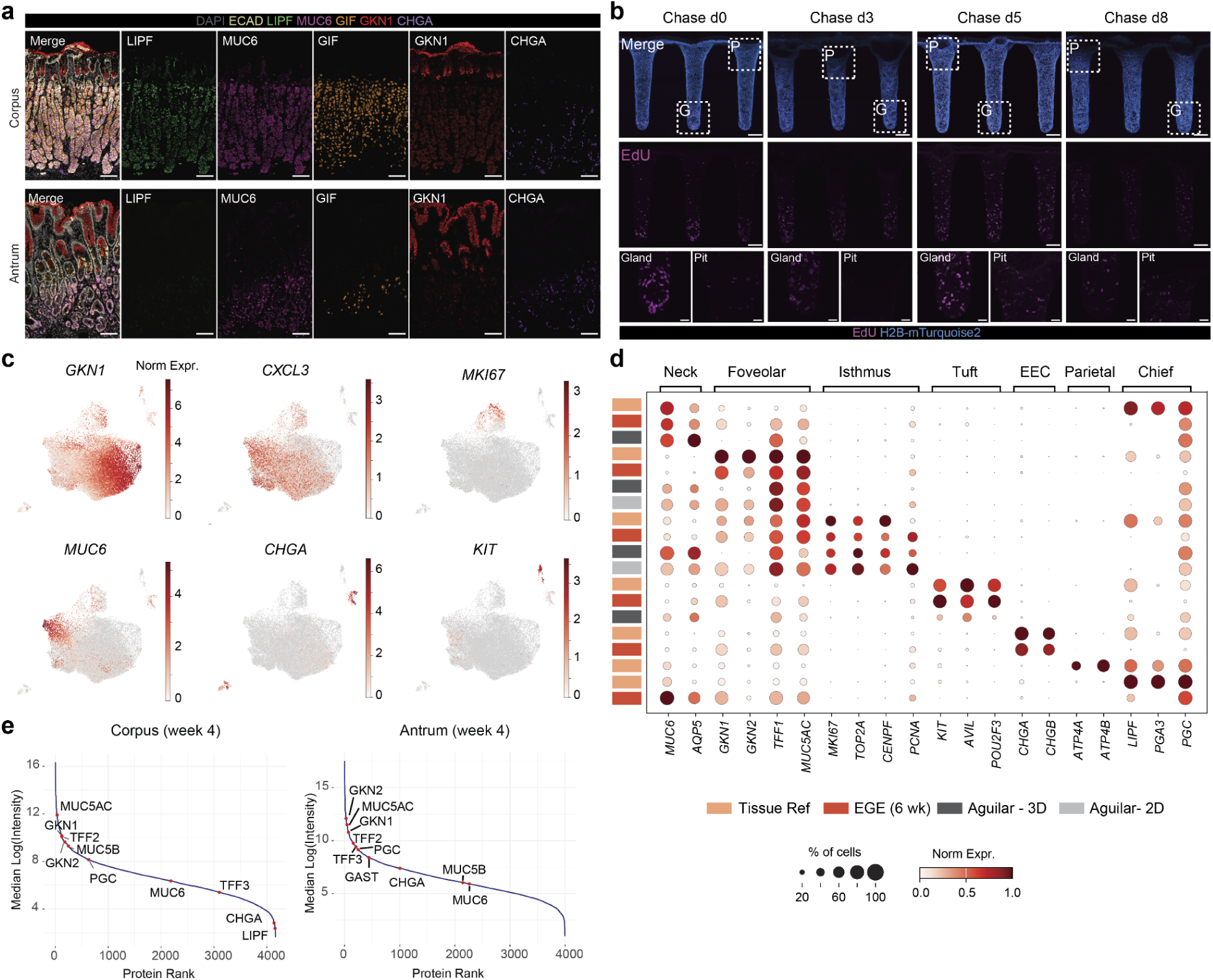
Characterization and benchmarking of EGE. **a**, Multiplexed immunofluorescence staining of adult human stomach antrum and corpus tissue for DAPI, E-cadherin (E-cad), LIPF, MUC6, GIF, GKN1, and CHGA, shown individually and as merged images. Scale bars: 100 μm. **b**, EdU pulse/chase of stomach cultures 3-4 weeks post seeding, showing EdU and the H2B-mTurquoise reporter along the gland–pit axis at four different chase time points. Scale bars overview: 100 μm, inserts: 25 μm. **c**, Feature plots displaying selected marker gene expression across the single-cell atlas. **d**, Dot plot of selected marker gene expression across adult stomach tissue, engineered EGE, Aguilar 2D monolayers, and Aguilar 3D organoids. **e**, Mass spectrometry analysis of the apical supernatant of corpus (Donor 1) and antrum (Donor 3) engineered tissues, with proteins ranked by abundance.

**Extended Data Fig. 3:**
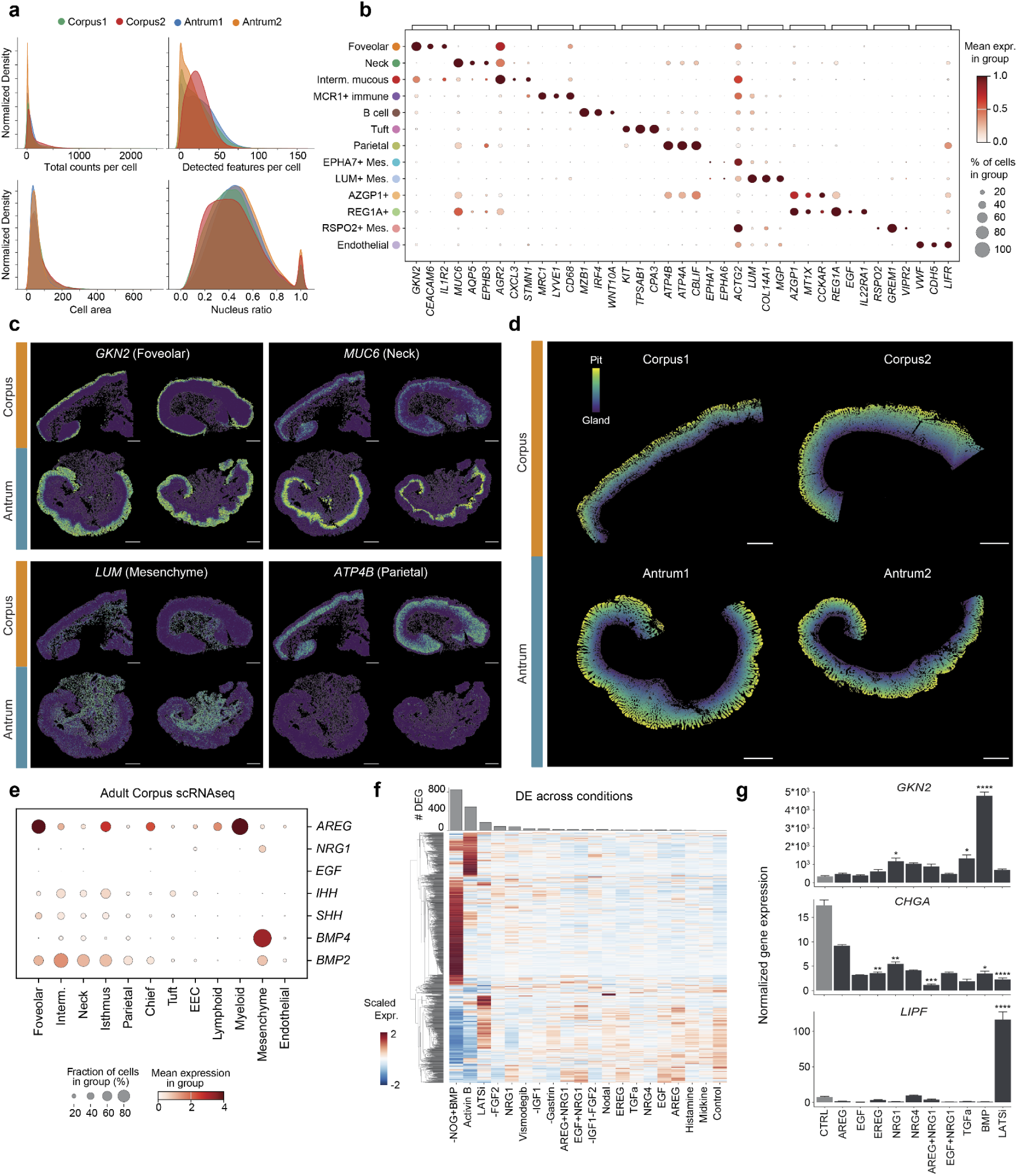
Quality control, cell-type annotation, and spatial patterning in Xenium datasets. **a**, Normalized density distributions of basic Xenium quality-control metrics, including total transcript counts per cell, number of detected features per cell, cell area, and nucleus-to-cell area ratio. **b**, Dot plot showing expression of selected marker genes used to annotate cell types and clusters in the merged Xenium dataset. **c**, Spatial feature plots of representative marker genes (*GKN2*, *MUC6*, *LUM*, *ATP4B*) shown for corpus and antrum tissue samples. Scale bars: 1 mm. **d**, Gradient masks to order cells along the gland-to-pit axis across all samples based on spatial position. Scale bars: 1 mm. **e**, Dot plot showing expression of foveolar-enriched genes in the single cell RNAseq tissue reference. **f**, Number of differentially expressed genes across 21 medium conditions, together with a heatmap showing scaled gene expression across treatments (n=3 replicate EGE cultured and sequenced independently per condition). **g**, Normalized gene expression of 4-week-old engineered stomach cultures upon 2-week treatment with the indicated molecules, showing *GKN2*, *CHGA* and *LIPF*.

**Extended Data Fig. 4:**
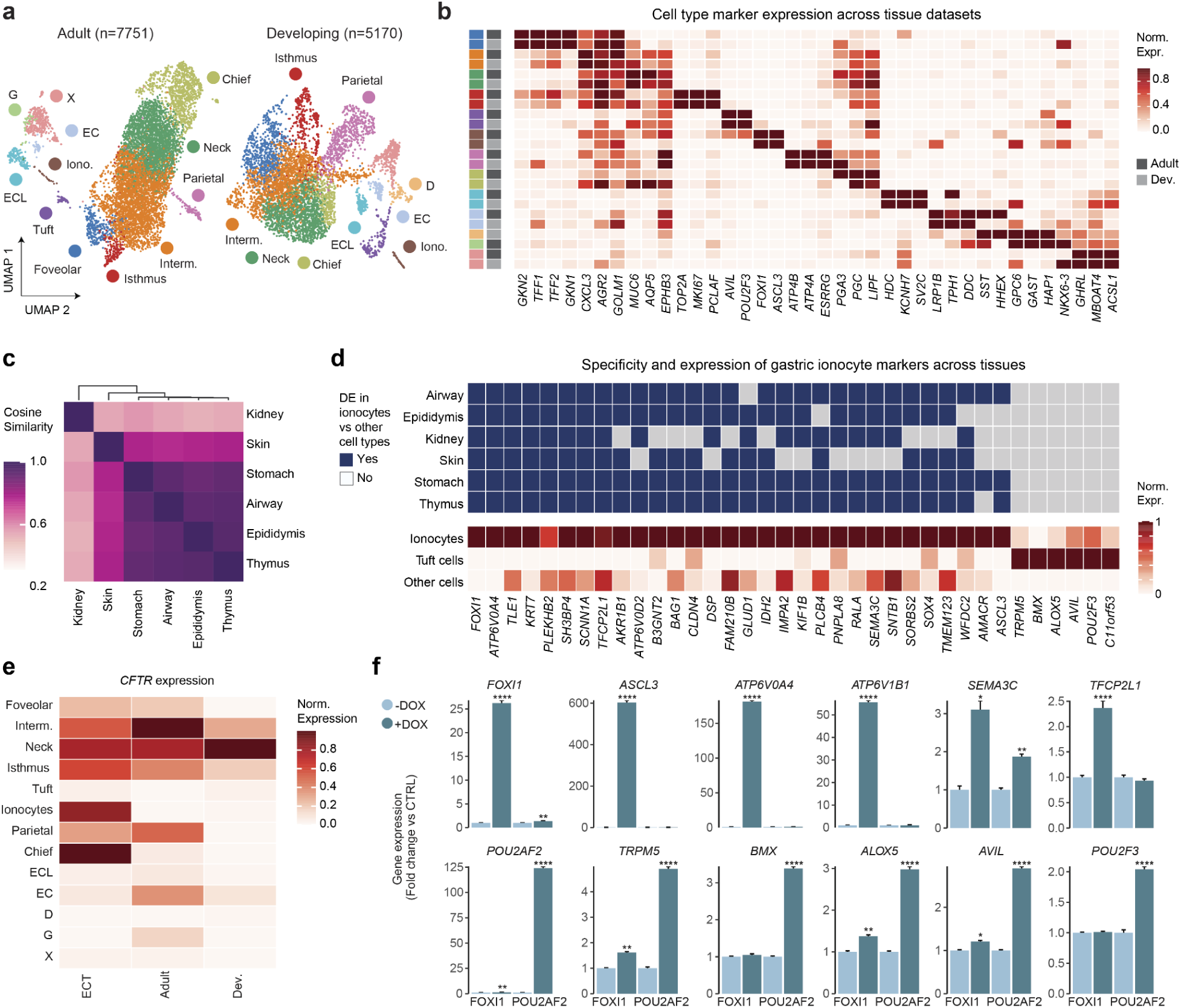
Characterization of gastric tuft cell and ionocyte lineages. **a**, UMAPs showing the annotation of adult and developing corpus tissue epithelial single cell RNAseq datasets. **b**, Heat map showing top DEGs for gastric lineages in the integrated gastric scRNAseq atlas from the developing and adult stomach. **c**, Cosine similarity plot showing the transcriptomic resemblance between ionocytes derived from the stomach, airway, epididymis, thymus, skin and kidney intercalated cells. **d**, Heatmap showing expression of gastric ionocyte signature genes compared to gastric tuft and other epithelial cells. The top panel indicates whether a gene is enriched in ionocytes in other tissues (blue) or not (gray). **e**, Heatmap showing normalized expression of *CFTR* across gastric lineages in developing tissue, adult tissue and EGE. **f**, Fold changes in expression of ionocyte-(top) and tuft cell-enriched (bottom) genes upon overexpression (DOX) of FOXI1 or POU2AF2 in corpus organoids versus controls (-DOX), measured by qPCR (n=3 samples per condition, each sample measured in technical triplicate).

**Extended Data Fig. 5:**
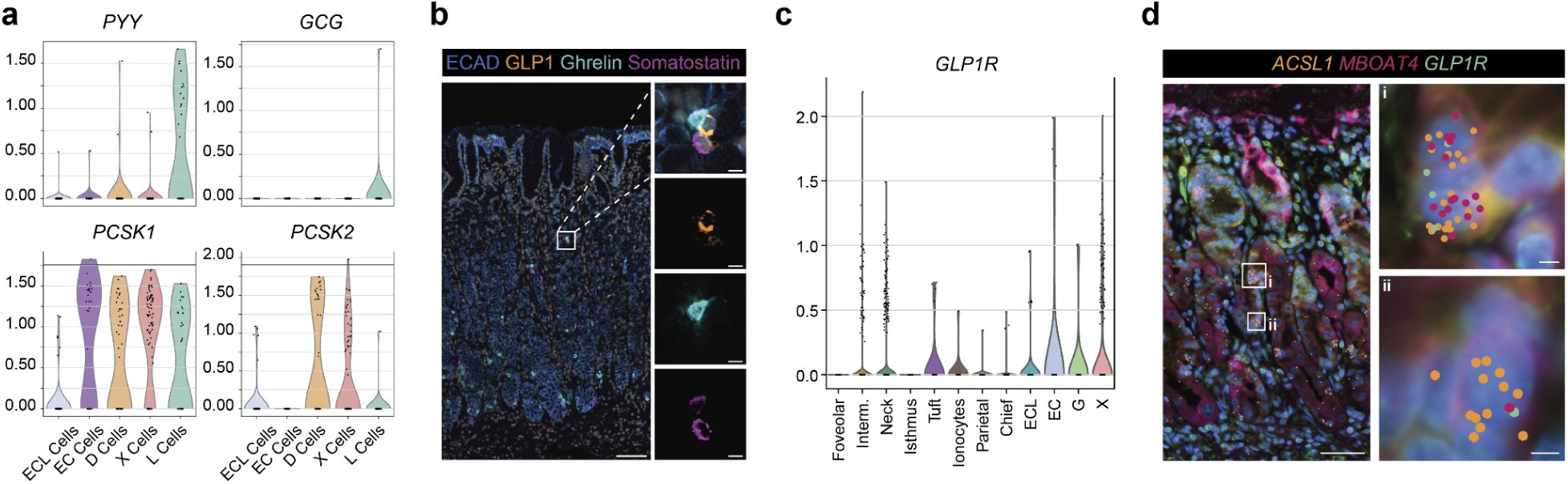
EGE contain rare GLP-1 producing gastric L cells. **a**, Violin plots showing expression of *PYY*, *GCG*, *PCSK1* and *PCSK2* expression in EGE EECs. **b**, Multiplexed immunofluorescent staining of hormones GLP-1, Ghrelin, Somatostatin and Histamine in corpus tissue (donor 2). Scale bar overview: 100 μm, inserts: 10 μm. **c**, Violin plots of the adult scRNAseq reference data set, showing expression of *GLP1R* across corpus epithelial lineages. **d**, IF image of corpus tissue (Donor 1) displaying Xenium data for *ACSL1*, *MBOAT4* (X cell markers) and *GLP1R*. Scale bar overview: 50 μm, inserts: 5 μm.

**Extended Data Fig. 6:**
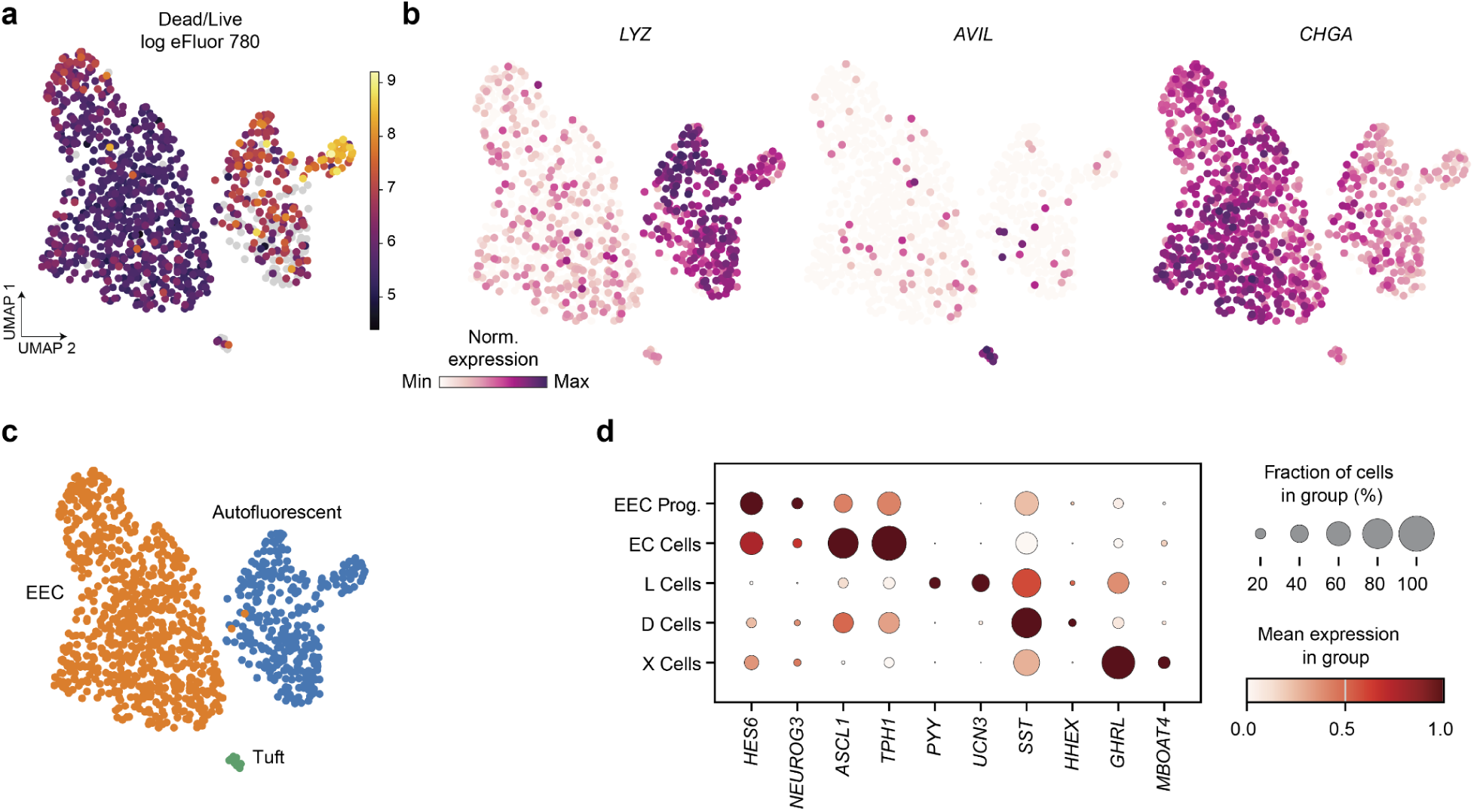
Quality control and annotation of NEUROG3-Chrono-derived scRNAseq data. **a**, Fluorescence intensities of eFluor 780 viability in cells processed for SORT-seq. **b**, Expression of *CHGA*, *LYZ* and *AVIL* in cells processed for SORT-seq. **c**, UMAP showing cells processed for SORT-seq, labeled as EECs, tuft cells or autofluorescent cells. **d**, Dot plot showing expression progenitor and mature EEC markers in EEC populations derived from NEUROG3-Chrono+ cells.

**Extended Data Fig. 7:**
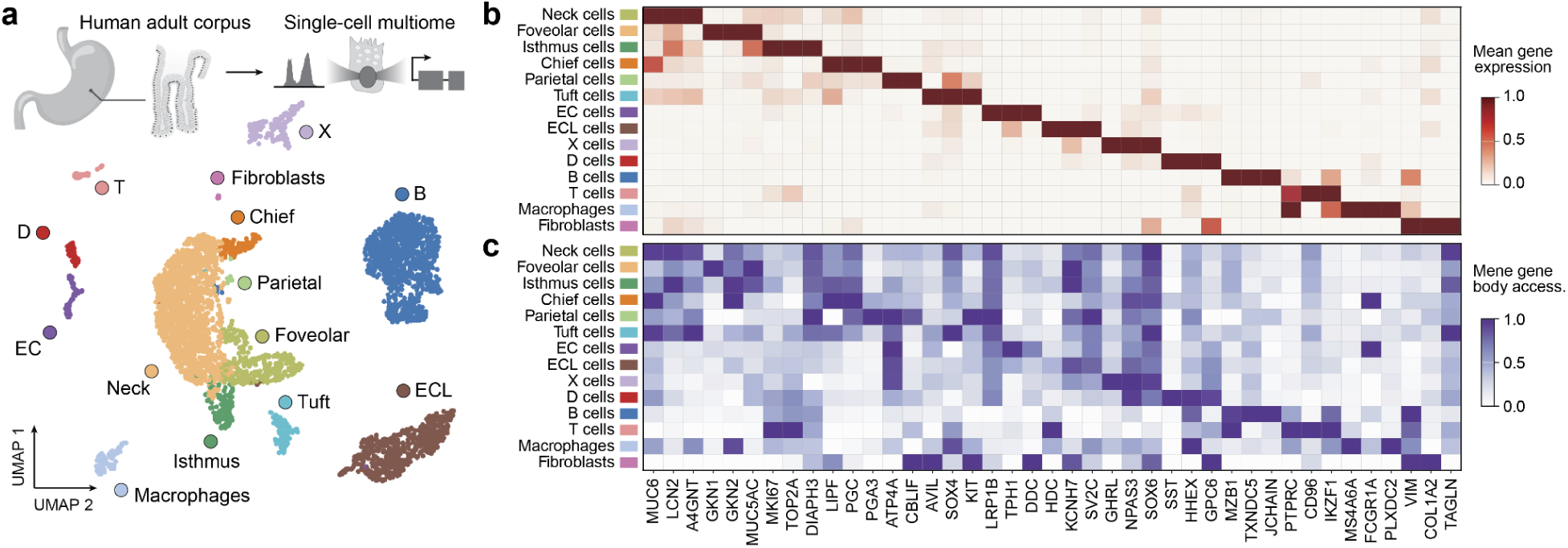
Multiomic atlas of the adult human stomach. **a**, Experimental design for adult human gastric corpus single-cell multiome profiling and UMAP embedding showing cellular heterogeneity in gene expression space. **b**, Cell type-specific marker gene expression and **c**, corresponding gene body chromatin accessibility across gastric cell populations.

**Extended Data Fig. 8:**
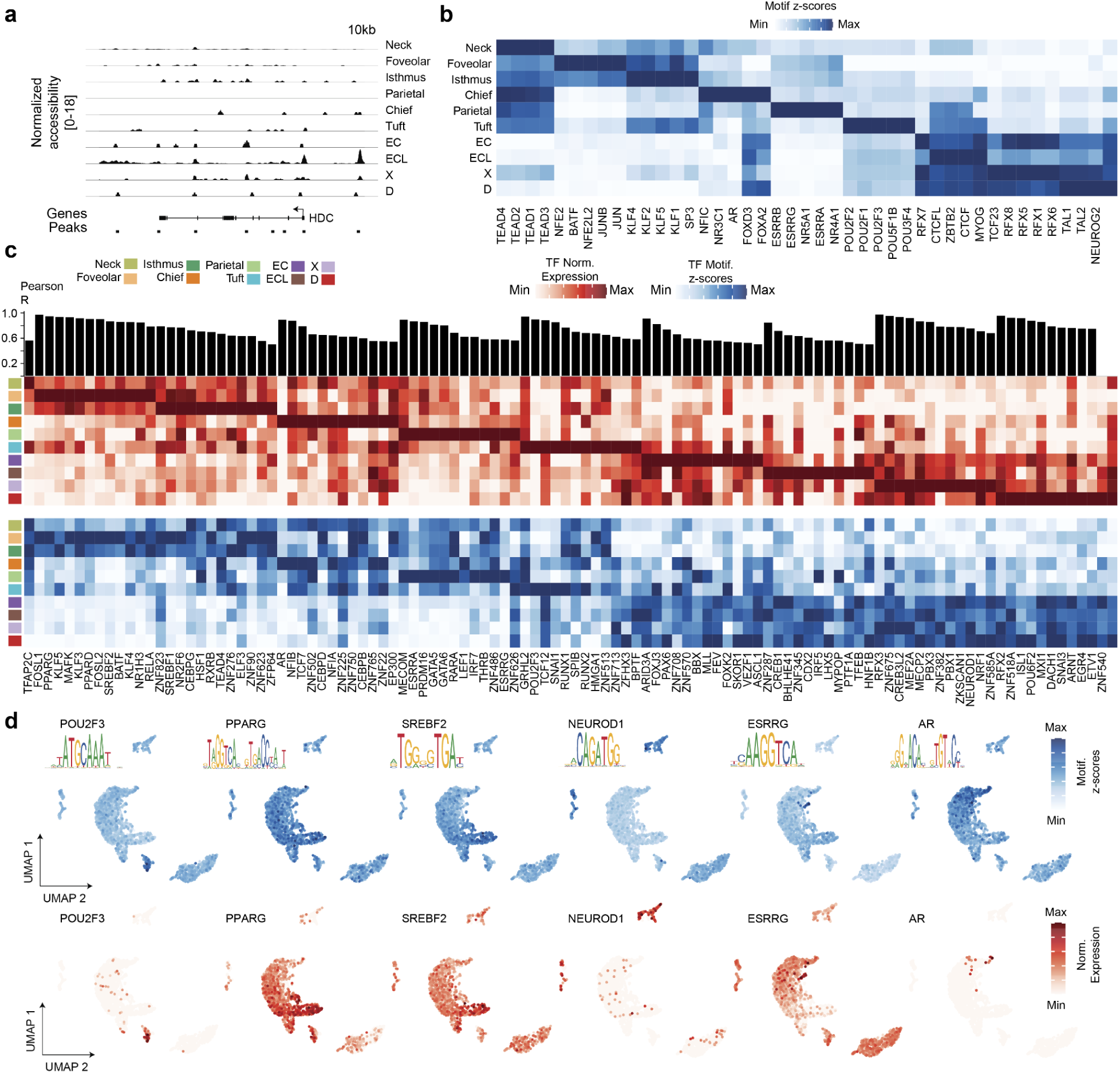
Chromatin accessibility landscapes and transcription factor enrichment across gastric cell types. **a**, Aggregate chromatin accessibility profiles at representative marker loci for enterochromaffin-like (ECL) cells (*HDC*). **b**, Top five enriched transcription factor motifs per cell type inferred using chromVAR. **c**, Top 12 transcription factors per cell type showing the highest correlation between transcription factor expression and motif accessibility across cell types. TFs with a lower correlation than 0.5 were removed. Heatmaps display normalized gene expression and motif *z*-scores, with transcription factors ordered by their maximal expression across cell types. **d**, UMAP embeddings colored by motif *z*-scores and normalized expression of selected epithelial cell type–enriched transcription factors.

**Extended Data Fig. 9:**
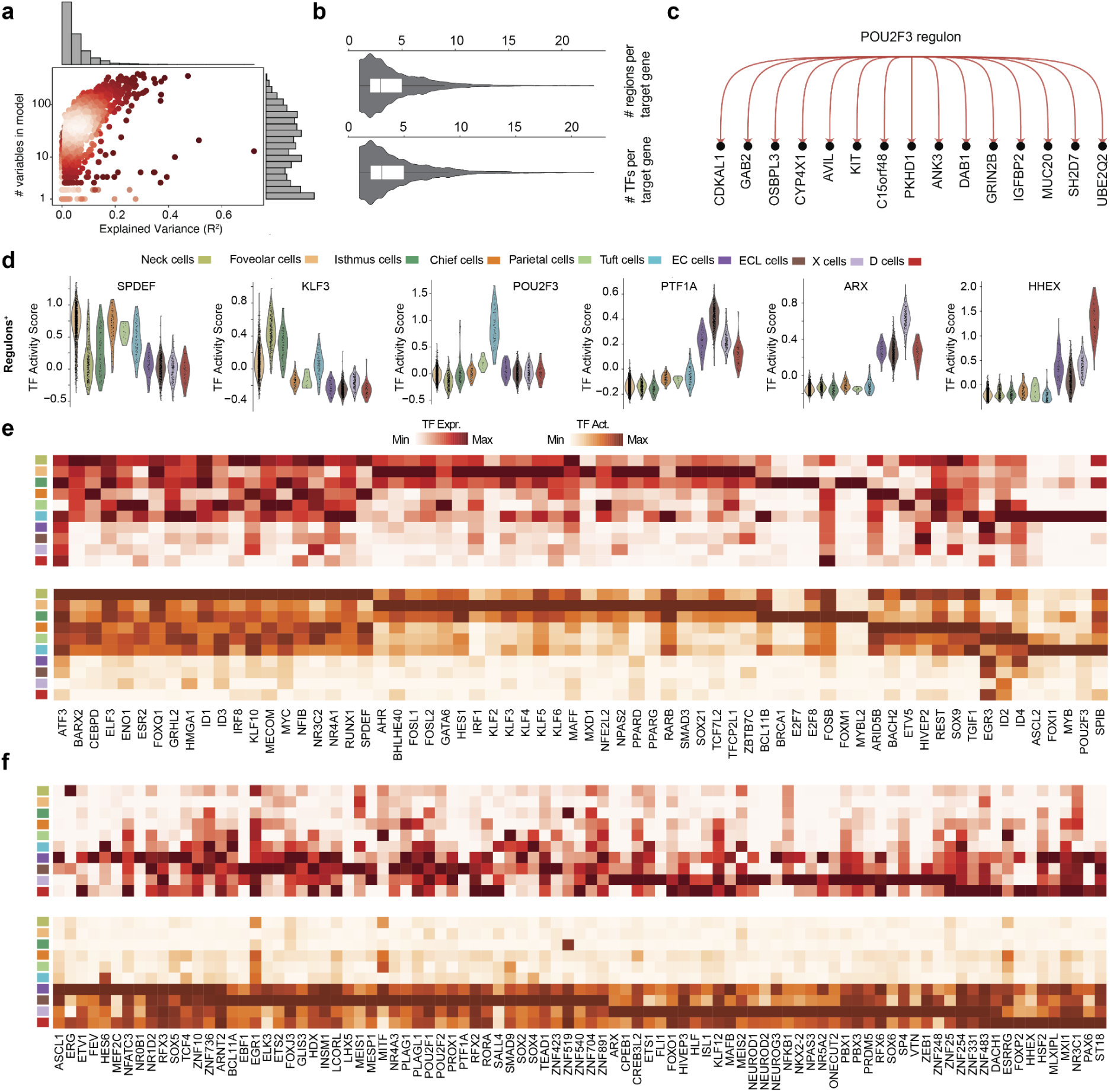
Pando infers cell-type enriched regulons and reveals TF-target gene interactions across epithelial lineages. **a**, Quality control scatter plot and accompanying histograms showing model performance of inferred Pando GRNs, displaying explained variance (Pearson R²) as a function of the number of predictor variables per model. **b**, Violin plots illustrating the distribution of the number of inferred regulatory regions per target gene and the number of transcription factors (TFs) associated with each target gene. **c**, Example regulon centered on *POU2F3*, as inferred by Pando, highlighting TF–target interactions. **d**, Violin plots displaying regulon activity scores (positive target genes) across cell types for SPDEF, KLF3, POU2F3, PTF1A, ARX, and HHEX. Activity scores were calculated from the positive target genes of each inferred TF regulon. **e**, Heatmap displaying normalized gene expression for each TF. Scores were computed per cell and averaged per cell type. **f**, Heatmap displaying regulon activity scores for each TF. Scores were computed per cell and averaged per cell type.

**Extended Data Table 1.** Media conditions screened in EGE.

| Condition | Details | Concentration |
|---|---|---|
| CTRL | Organoid expansion medium | NA |
| -NOG+BMP | Noggin removed, BMP2 and BMP4 supplemented | Both 50 ng/mL |
| Activin B | Activin B supplemented | 50 ng/mL |
| LATSi | LATSi supplemented | 5 µM |
| NRG1 | NRG1 supplemented | 50 ng/mL |
| -FGF2 | FGF2 removed | NA |
| Vismodegib | Vismodegib (Hedgehog inhibitor) | 1 µM |
| -IGF1 | IGF1 removed | NA |
| -Gastrin | Gastrin removed | NA |
| AREG + NRG1 | AREG and NRG1 supplemented | Both 50 ng/mL |
| -IGF1 -FGF2 | IGF1 and FGF2 removed | NA |
| EGF+NRG1 | EGF and NRG1 supplemented | Both 50 ng/mL |
| TGFa | TGF-alpha supplemented | 50 ng/mL |
| EREG | EREG supplemented | 50 ng/mL |
| NRG4 | NRG4 supplemented | 50 ng/mL |
| Nodal | Nodal supplemented | 50 ng/mL |
| EGF | EGF supplemented | 50 ng/mL |
| AREG | AREG supplemented | 50 ng/mL |
| NRG1 (low) | NRG1 supplemented (low concentration) | 10 ng/mL |
| Histamine | Histamine supplemented | 50 ng/mL |
| Midkine | Midkine supplemented | 50 ng/mL |

**Extended Data Table 2.** Medium components used in this study.

| **Component** | **Vendor** | **Cat** |
|---|---|---|
| Advanced DMEM/F12 | Thermo Fisher Scientific | Cat #12634-010 |
| Dulbecco’s Phosphate-buffered saline (DPBS) | Thermo Fisher Scientific | Cat #J67670.AP |
| B-27 Supplement minus Vitamin A | Thermo Fisher Scientific | Cat #17504044 |
| GlutaMAX | Thermo Fisher Scientific | Cat #35050061 |
| HEPES | Thermo Fisher Scientific | Cat #15630056 |
| Penicillin-Streptomycin | Thermo Fisher Scientific | Cat #15140122 |
| Primocin | Invivogen | Cat #ant-pm-2 |
| Matrigel Growth Factor Reduced | Corning | Cat #356231 |
| Rspo3-Fc Fusion Protein conditioned media | ImmunoPrecise Antibodies | Cat #R001 |
| Noggin-Fc Fusion Protein conditioned media | ImmunoPrecise Antibodies | Cat #N002 |
| WNT Surrogate-FC Fusion Protein | ImmunoPrecise Antibodies | Cat #N001 |
| Human Gastrin I | MedChemExpress | Cat #HY-P1097 |
| A83-01 | Tocris | Cat #2939 |
| Human FGF-10 | Peprotech | Cat #100-26 |
| Human FGF-basic (FGF-2/bFGF) | Thermo Fisher Scientific | Cat #100-18B |
| Y-27632 dihydrochloride | Tocris | Cat #1254 |
| Forskolin | Tocris | Cat #1099 |
| IBMX | MedChemExpress | Cat #HY-12318 |
| Recombinant Human Amphiregulin | Peprotech | Cat #100-55B |
| Neuregulin-4 Protein, Human | MedChemExpress | Cat #HY-P76524 |
| Human BMP-2 Recombinant Protein | Thermo Fisher Scientific | Cat #120-02 |
| Human BMP-4 Recombinant Protein | Thermo Fisher Scientific | Cat #120-05ET |
| CHIR 99021 | Tocris | Cat #4423 |
| Human EGF, Animal-Free Recombinant Protein | Thermo Fisher Scientific | Cat #AF-100-15 |
| Recombinant Human Neuregulin-1 | R&D Systems | Cat #5898-NR |
| Recombinant Human IGF-I | Biolegend | Cat #590908 |
| Human IGF-II Recombinant Protein | Thermo Fisher Scientific | Cat #100-12 |
| Recombinant Human Activin B Protein, CF | R&D Systems | Cat #11517-AB |
| TRULI (LATS inhibitor) | MedChemExpress | Cat #HY-138489 |
| Vismodegib | MedchemExpress | Cat #HY-10440 |
| Recombinant Human Nodal Protein | R&D Systems | Cat #3218-ND-025 |
| Human TGF-alpha Recombinant Protein | Peprotech | Cat #100-16A |
| Human Sonic Hedgehog C25II Recombinant Protein | Thermo Fisher Scientific | Cat #100-45 |
| Recombinant Human Epiregulin Protein | R&D Systems | Cat #1195-EP-025 |
| Histamine | MedchemExpress | Cat #HY-B1204 |
| Recombinant Human Midkine | Peprotech | Cat #450-16 |
| Human Indian Hedgehog | MedchemExpress | Cat #HY-P7204 |

**Extended Data Table 3.** Antibodies used in this study.

| Antibody | Vendor | Cat |
|---|---|---|
| anti-CHGA | Abcam | #ab254557 |
| anti-GIF | Atlas Antibodies | #HPA039908 |
| anti-MUC6 | Abcam | #ab223846 |
| anti-LIPF | Atlas Antibodies | #HPA045930 |
| anti-GKN1 | Atlas Antibodies | #HPA047684 |
| anti-Gastrin | Thermo Fisher Scientific | #MA5-50699 |
| anti-Histamine | Antibodies Online | #ABIN7427816 |
| anti-GLP-1 | Abcam | Cat: #ab23468 |
| anti-Ghrelin | Abcam | Cat: #ab209790 |
| anti-Somatostatin | Abcam | Cat: #ab111912 |
| anti-Serotonin | Abcam | Cat: #ab66047 |
| anti-ATP6V1B1 | Atlas Antibodies | Cat: #HPA031847 |
| anti-CD117 (c-kit) | BioLegend | Cat: #375302 |
| anti-E-cadherin | Ventana | Cat: #05905290001 |
| anti-Rabbit HRP | OmniMap | #760-4311 |
| anti-Mouse HRP | OmniMap | #760-4310 |
| APC anti-CD326 (Ep-CAM) | BioLegend | Cat: #324208 |
| FITC anti-CD326 (Ep-CAM) | BioLegend | Cat: #324204 |
| PE/Cyanine7 anti-human CD117 (c-kit) | BioLegend | Cat: #323414 |
| APC anti-human CD200 | BioLegend | Cat: #329208 |
| Purified anti-human CD117 (c-kit) | BioLegend | Cat: #375302 |
| Phalloidin Alexa Fluor Plus 750 | Thermo Fisher Scientific | Cat: #A30105 |
| Donkey anti-Rabbit Alexa Fluor Plus 555 | Thermo Fisher Scientific | Cat: #A32794 |
| Donkey anti-Mouse Alexa Fluor Plus 488 | Thermo Fisher Scientific | Cat: #A21202 |
| Donkey anti-Rat Alexa Fluor Plus 647 | Thermo Fisher Scientific | Cat: #A48272TR |
| Donkey anti-Goat DyeLight 755 | Thermo Fisher Scientific | Cat: #SA5-10091l |

## References

1. Willet, S. G. & Mills, J. C. Stomach Organ and Cell Lineage Differentiation: from Embryogenesis to Adult Homeostasis. Cell Mol Gastroenterol Hepatol 2, 546–559 (2016).

2. Bartfeld, S. & Koo, B.-K. Adult gastric stem cells and their niches. Wiley Interdiscip Rev Dev Biol 6, (2017).

3. Kim, T.-H. & Shivdasani, R. A. Stomach development, stem cells and disease. Development 143, 554–565 (2016).

4. Busslinger, G. A. et al. Human gastrointestinal epithelia of the esophagus, stomach, and duodenum resolved at single-cell resolution. Cell Rep 34, 108819 (2021).

5. Choi, E. et al. Cell lineage distribution atlas of the human stomach reveals heterogeneous gland populations in the gastric antrum. Gut 63, 1711–1720 (2014).

6. Tan, S. H. et al. AQP5 enriches for stem cells and cancer origins in the distal stomach. Nature 578, 437–443 (2020).

7. Lim, H. Y. G. et al. AQP5: A functional gastric cancer stem cell marker in mouse and human tumors. Science (2025).

8. Huh, W. J. et al. Tamoxifen induces rapid, reversible atrophy, and metaplasia in mouse stomach. Gastroenterology 142, 21–24.e7 (2012).

9. Mills, J. C. & Goldenring, J. R. Metaplasia in the Stomach Arises From Gastric Chief Cells. Cell Mol Gastroenterol Hepatol 4, 85–88 (2017).

10. Nam, K. T. et al. Mature chief cells are cryptic progenitors for metaplasia in the stomach. Gastroenterology 139, 2028–2037.e9 (2010).

11. Bartfeld, S. et al. In vitro expansion of human gastric epithelial stem cells and their responses to bacterial infection. Gastroenterology 148, 126–136.e6 (2015).

12. Barker, N. et al. Lgr5(+ve) stem cells drive self-renewal in the stomach and build long-lived gastric units in vitro. Cell Stem Cell 6, 25–36 (2010).

13. Eicher, A. K. et al. Functional human gastrointestinal organoids can be engineered from three primary germ layers derived separately from pluripotent stem cells. Cell Stem Cell 29, 36–51.e6 (2022).

14. Broda, T. R., McCracken, K. W. & Wells, J. M. Generation of human antral and fundic gastric organoids from pluripotent stem cells. Nature Protocols 14, 28–50 (2018).

15. McCracken, K. W. et al. Modelling human development and disease in pluripotent stem-cell-derived gastric organoids. Nature 516, 400–404 (2014).

16. Plasschaert, L. W. et al. A single-cell atlas of the airway epithelium reveals the CFTR-rich pulmonary ionocyte. Nature 560, 377–381 (2018).

17. Montoro, D. T. et al. A revised airway epithelial hierarchy includes CFTR-expressing ionocytes. Nature 560, 319–324 (2018).

18. Yuan, F. et al. Transgenic ferret models define pulmonary ionocyte diversity and function. Nature 621, 857–867 (2023).

19. Vidarsson, H. et al. The forkhead transcription factor Foxi1 is a master regulator of vacuolar H-ATPase proton pump subunits in the inner ear, kidney and epididymis. PLoS One 4, e4471 (2009).

20. Quigley, I. K., Stubbs, J. L. & Kintner, C. Specification of ion transport cells in the Xenopus larval skin. Development 138, 705–714 (2011).

21. Meng, Y. H. Intracellular PH Regulation in H+-ATPase-Rich Ionocytes in Zebrafish Larvae Using in Vivo Ratiometric Imaging. (2017).

22. Aguilar, C. et al. Helicobacter pylori shows tropism to gastric differentiated pit cells dependent on urea chemotaxis. Nat Commun 13, 5878 (2022).

23. Bollen, Y. et al. Efficient and error-free fluorescent gene tagging in human organoids without double-strand DNA cleavage. PLoS Biol 20, e3001527 (2022).

24. Serra, D. et al. Self-organization and symmetry breaking in intestinal organoid development. Nature 569, 66–72 (2019).

25. Schwayer, C. et al. Cell heterogeneity and fate bistability drive tissue patterning during intestinal regeneration. bioRxiv (2025) doi:10.1101/2025.01.14.632683.

26. Fujii, M. et al. Human Intestinal Organoids Maintain Self-Renewal Capacity and Cellular Diversity in Niche-Inspired Culture Condition. Cell Stem Cell 23, 787–793.e6 (2018).

27. Gjorevski, N. et al. Tissue geometry drives deterministic organoid patterning. Science 375, eaaw9021 (2022).

28. Mitrofanova, O. et al. Bioengineered human colon organoids with in vivo-like cellular complexity and function. Cell Stem Cell 31, 1175–1186.e7 (2024).

29. Nikolaev, M. et al. Homeostatic mini-intestines through scaffold-guided organoid morphogenesis. Nature 585, 574–578 (2020).

30. López-Sandoval, R., et al. Modeling host-microbe interactions in immunocompetent engineered human gut tissues. bioRxiv (2025) doi:10.1101/2025.05.02.651468.

31. Lee, S.-H. et al. Apposition of Fibroblasts With Metaplastic Gastric Cells Promotes Dysplastic Transition. Gastroenterology 165, 374–390 (2023).

32. Hong, F. et al. The critical role of BMP signaling in gastric epithelial cell differentiation revealed by organoids. Cell Regen. (Lond.) 14, 18 (2025).

33. Lo, H. G. et al. A single transcription factor is sufficient to induce and maintain secretory cell architecture. Genes & development 31, (2017).

34. Ballantyne, G. H. et al. Vasoactive intestinal peptide augmentation of cholecystokinin-8-stimulated pepsinogen secretion: evidence for dual modulation of chief cell function. The Journal of surgical research 40, (1986).

35. Raffaniello, R. D. & Raufman, J. P. Pepsinogen secretion from streptolysin O-permeabilized chief cells from guinea pig stomach. Am J Physiol 263, G452–9 (1992).

36. Modlin, I. M. et al. Forskolin (cyclic adenosine monophosphate)-dependent protein phosphorylation in isolated gastric glands. Arch Surg 121, 330–337 (1986).

37. Janesick, A. et al. High resolution mapping of the tumor microenvironment using integrated single-cell, spatial and in situ analysis. Nat Commun 14, 8353 (2023).

38. Wölffling, S., et al. EGF and BMPs Govern Differentiation and Patterning in Human Gastric Glands. Gastroenterology 161, 623–636.e16 (2021).

39. Konstantinou, D., Bertaux-Skeirik, N. & Zavros, Y. Hedgehog signaling in the stomach. Curr Opin Pharmacol 31, 76–82 (2016).

40. Kapalczynska, M. et al. BMP feed-forward loop promotes terminal differentiation in gastric glands and is interrupted by H. pylori-driven inflammation. Nature Communications 13, 1577 (2022).

41. van den Brink, G. R., et al. Sonic hedgehog regulates gastric gland morphogenesis in man and mouse. Gastroenterology 121, 317–328 (2001).

42. Fukaya, M. et al. Hedgehog signal activation in gastric pit cell and in diffuse-type gastric cancer. Gastroenterology 131, 14–29 (2006).

43. Walton, K. D. et al. Villification in the mouse: Bmp signals control intestinal villus patterning. Development 143, 427–436 (2016).

44. Walton, K. D. & Gumucio, D. L. Hedgehog Signaling in Intestinal Development and Homeostasis. Annu Rev Physiol 83, 359–380 (2021).

45. Basak, O. et al. Induced Quiescence of Lgr5+ Stem Cells in Intestinal Organoids Enables Differentiation of Hormone-Producing Enteroendocrine Cells. Cell Stem Cell 20, 177–190.e4 (2017).

46. Kim, D.-H. et al. A type 2 immune circuit in the stomach controls mammalian adaptation to dietary chitin. Science 381, 1092–1098 (2023).

47. Haber, A. L. et al. A single-cell survey of the small intestinal epithelium. Nature 551, 333–339 (2017).

48. Huang, L. et al. Tuft cells act as regenerative stem cells in the human intestine. Nature 634, 929–935 (2024).

49. Buissant des Amorie, J. R., et al. Intestinal tuft cell subtypes represent successive stages of maturation driven by crypt-villus signaling gradients. Nat Commun 16, 6765 (2025).

50. Beumer, J. et al. Description and functional validation of human enteroendocrine cell sensors. Science 386, 341–348 (2024).

51. Pou Casellas, C., Pleguezuelos-Manzano, C., Rookmaaker, M. B., Verhaar, M. C. & Clevers, H. Transcriptomic profile comparison reveals conservation of ionocytes across multiple organs. Sci Rep 13, 3516 (2023).

52. Bautista, J. L. et al. Single-cell transcriptional profiling of human thymic stroma uncovers novel cellular heterogeneity in the thymic medulla. Nat Commun 12, 1096 (2021).

53. Leir, S.-H., Yin, S., Kerschner, J. L., Cosme, W. & Harris, A. An atlas of human proximal epididymis reveals cell-specific functions and distinct roles for CFTR. Life Sci Alliance 3, (2020).

54. Ravindra, N. G. et al. Single-cell longitudinal analysis of SARS-CoV-2 infection in human airway epithelium identifies target cells, alterations in gene expression, and cell state changes. PLoS Biol 19, e3001143 (2021).

55. Travaglini, K. J. et al. A molecular cell atlas of the human lung from single-cell RNA sequencing. Nature 587, 619–625 (2020).

56. He, H. et al. Single-cell transcriptome analysis of human skin identifies novel fibroblast subpopulation and enrichment of immune subsets in atopic dermatitis. J Allergy Clin Immunol 145, 1615–1628 (2020).

57. Muto, Y. et al. Single cell transcriptional and chromatin accessibility profiling redefine cellular heterogeneity in the adult human kidney. Nat Commun 12, 2190 (2021).

58. Huang, Y. H. et al. POU2F3 is a master regulator of a tuft cell-like variant of small cell lung cancer. Genes & development 32, (2018).

59. Wu, X. S. et al. OCA-T1 and OCA-T2 are coactivators of POU2F3 in the tuft cell lineage. Nature 607, 169–175 (2022).

60. van der Meulen, T. et al. Urocortin3 mediates somatostatin-dependent negative feedback control of insulin secretion. Nature medicine 21, 769 (2015).

61. Ribeiro-Parenti, L. et al. Bariatric surgery induces a new gastric mucosa phenotype with increased functional glucagon-like peptide-1 expressing cells. Nature Communications 12, 110 (2021).

62. Müller, T. D., Finan, B., Clemmensen, C., DiMarchi, R. D. & Tschöp, M. H. The New Biology and Pharmacology of Glucagon. Physiol Rev 97, 721–766 (2017).

63. Roberts, G. P. et al. Comparison of Human and Murine Enteroendocrine Cells by Transcriptomic and Peptidomic Profiling. Diabetes 68, 1062–1072 (2019).

64. Gehart, H. et al. Identification of Enteroendocrine Regulators by Real-Time Single-Cell Differentiation Mapping. Cell 176, 1158–1173.e16 (2019).

65. Muraro, M. J. et al. A Single-Cell Transcriptome Atlas of the Human Pancreas. Cell Syst 3, 385–394.e3 (2016).

66. Singh, P. N. P. et al. Transcription factor dynamics, oscillation, and functions in human enteroendocrine cell differentiation. Cell Stem Cell 31, 1038–1057.e11 (2024).

67. Zhang, J., McKenna, L. B., Bogue, C. W. & Kaestner, K. H. The diabetes gene Hhex maintains δ-cell differentiation and islet function. Genes Dev 28, 829–834 (2014).

68. Terry, N. A., Walp, E. R., Lee, R. A., Kaestner, K. H. & May, C. L. Impaired enteroendocrine development in intestinal-specific Islet1 mouse mutants causes impaired glucose homeostasis. Am J Physiol Gastrointest Liver Physiol 307, G979–91 (2014).

69. Fleck, J. S. et al. Inferring and perturbing cell fate regulomes in human brain organoids. Nature 621, 365–372 (2023).

70. Katz, J. P. et al. The zinc-finger transcription factor Klf4 is required for terminal differentiation of goblet cells in the colon. Development 129, 2619–2628 (2002).

71. Verzi, M. P., Khan, A. H., Ito, S. & Shivdasani, R. A. Transcription factor foxq1 controls mucin gene expression and granule content in mouse stomach surface mucous cells. Gastroenterology 135, 591–600 (2008).

72. Willet, S. G. et al. SOX9 Governs Gastric Mucous Neck Cell Identity and Is Required for Injury-Induced Metaplasia. Cell Mol Gastroenterol Hepatol 16, 325–339 (2023).

73. Horst, D. et al. Requirement of the epithelium-specific Ets transcription factor Spdef for mucous gland cell function in the gastric antrum. J Biol Chem 285, 35047–35055 (2010).

74. Gracz, A. D. et al. Sox4 Promotes Atoh1-Independent Intestinal Secretory Differentiation Toward Tuft and Enteroendocrine Fates. Gastroenterology 155, 1508–1523.e10 (2018).

75. Hori, K. et al. A nonclassical bHLH–Rbpj transcription factor complex is required for specification of GABAergic neurons independent of Notch signaling. Genes Dev. 22, 166–178 (2008).

76. Adkins-Threats, M. et al. Metabolic regulator ERRγ governs gastric stem cell differentiation into acid-secreting parietal cells. Cell Stem Cell 31, 886–903.e8 (2024).

77. Jenny, M. et al. Neurogenin3 is differentially required for endocrine cell fate specification in the intestinal and gastric epithelium. EMBO J 21, 6338–6347 (2002).

78. Hibdon, E. S., Keeley, T. M., Merchant, J. L. & Samuelson, L. C. The bHLH transcription factor ASCL1 promotes differentiation of endocrine cells in the stomach and is regulated by Notch signaling. Am J Physiol Gastrointest Liver Physiol 325, G458–G470 (2023).

79. Flasse, L. C. et al. The bHLH transcription factor Ascl1a is essential for the specification of the intestinal secretory cells and mediates Notch signaling in the zebrafish intestine. Dev Biol 376, 187–197 (2013).

80. Kokubu, H., Ohtsuka, T. & Kageyama, R. Mash1 is required for neuroendocrine cell development in the glandular stomach. Genes Cells 13, 41–51 (2008).

81. Sesotyosari, S. L. et al. The long-term survival of enteroendocrine cells depends on their subtype and is linked to peripheral sensory innervation. Dev Growth Differ 67, 205–214 (2025).

82. Broide, E. et al. GLP-1 receptor is expressed in human stomach mucosa: analysis of its cellular association and distribution within gastric glands. J Histochem Cytochem 61, 649–658 (2013).

83. Grunddal, K. V. et al. Expression Profile of the GLP-1 Receptor in the Gastrointestinal Tract and Pancreas in Adult Female Mice. Endocrinology 163, (2022).

84. Schmidtler, J. et al. Rat parietal cell receptors for GLP-1-(7-36) amide: northern blot, cross-linking, and radioligand binding. Am J Physiol 267, G423–32 (1994).

85. Pleguezuelos-Manzano, C. et al. Establishment and Culture of Human Intestinal Organoids Derived from Adult Stem Cells. Current Protocols in Immunology 130, e106 (2020).

86. Fujii, M., Matano, M., Nanki, K. & Sato, T. Efficient genetic engineering of human intestinal organoids using electroporation. Nat Protoc 10, 1474–1485 (2015).

87. Leutert, M., Rodríguez-Mias, R. A., Fukuda, N. K. & Villén, J. R2-P2 rapid-robotic phosphoproteomics enables multidimensional cell signaling studies. Mol Syst Biol 15, e9021 (2019).

88. Kohler, D. et al. MSstats Version 4.0: Statistical Analyses of Quantitative Mass Spectrometry-Based Proteomic Experiments with Chromatography-Based Quantification at Scale. J Proteome Res 22, 1466–1482 (2023).

89. Lopez, R., Regier, J., Cole, M. B., Jordan, M. I. & Yosef, N. Deep generative modeling for single-cell transcriptomics. Nat. Methods 15, (2018).

90. Xu, C. et al. Probabilistic harmonization and annotation of single-cell transcriptomics data with deep generative models. Mol. Syst. Biol. 17, e9620 (2021).

91. Yanai, I. et al. Genome-wide midrange transcription profiles reveal expression level relationships in human tissue specification. Bioinformatics 21, 650–659 (2005).

92. Polański, K. et al. BBKNN: fast batch alignment of single cell transcriptomes. Bioinformatics 36, 964–965 (2020).

93. Dobin, A. et al. STAR: ultrafast universal RNA-seq aligner. Bioinformatics 29, 15–21 (2013).

94. Granja, J. M. et al. ArchR is a scalable software package for integrative single-cell chromatin accessibility analysis. Nat. Genet. 53, 403–411 (2021).

95. Schep, A. N., Wu, B., Buenrostro, J. D. & Greenleaf, W. J. chromVAR: inferring transcription-factor-associated accessibility from single-cell epigenomic data. Nat. Methods 14, 975–978 (2017).

96. Stuart, T., Srivastava, A., Madad, S., Lareau, C. A. & Satija, R. Single-cell chromatin state analysis with Signac. Nat. Methods 18, 1333–1341 (2021).

97. Benjamini, Y. & Hochberg, Y. Controlling the false discovery rate: A practical and powerful approach to multiple testing. J. R. Stat. Soc. Series B Stat. Methodol. 57, 289–300 (1995).

