## Supplementary Information for "A long-lived human gastric tissue model defines regulators of lineage maturation"

This file contains:

Supplementary Figure 1

Description of additional Supplementary Files

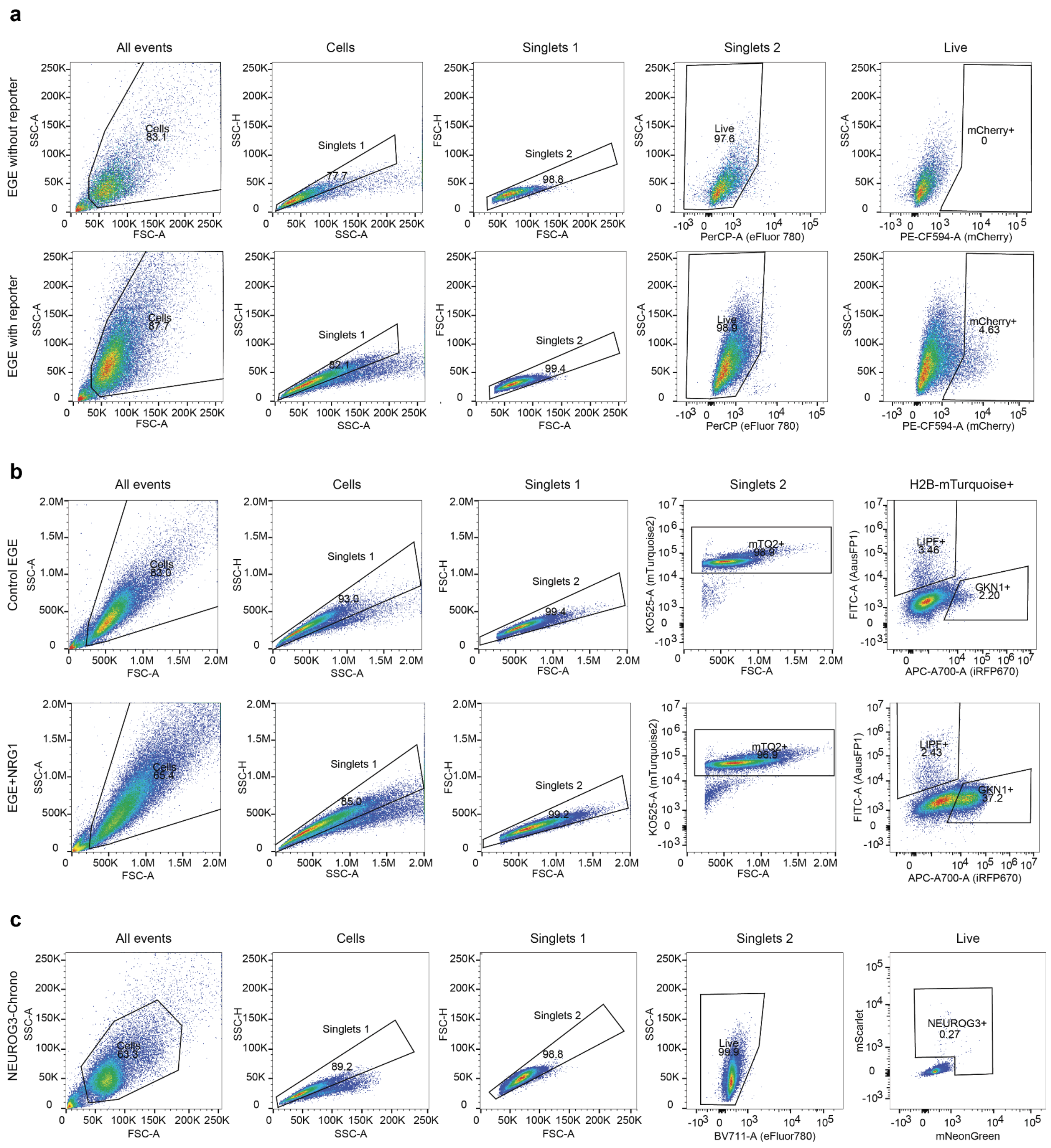


**Supplementary Fig. 1: Flow cytometry and FACs gating strategies.**

**a**, Gating strategy for FACs of EGE expressing ATP4A-mCherry and a control without reporter. **b**, Gating strategy for flow cytometry of EGE expressing GKN1-iRFP670 and LIPF-AausFP1 in control medium or supplemented with NRG1. **c**, Gating strategy for FACs of NEUROG3-Chrono+ cells.

**Description of additional Supplementary Files**

**Supplementary Video 1**
Homeostatic long-term culture of engineered gastric epithelium for over 6 months
https://youtu.be/jEK1Xw1Ixxw

**Supplementary Video 2**
Chief cell patterning at the base of glands in 1-week-old engineered gastric epithelium
https://youtu.be/YtROAQBIDCM

**Supplementary Video 3**
Parietal cells in 6-week-old engineered gastric epithelium
https://youtu.be/0g5QA0l5Z2I

**Supplementary Video 4**
Mature and functional gastric enteroendocrine cells in 4-week-old engineered epithelium
https://youtu.be/gblwzfVvNRg

**Supplementary Video 5**
Macro-scale engineered corpus epithelium
https://youtu.be/0mRH2XVtTfE

**Supplementary Table 1.**
Proteomics measurements

**Supplementary Table 2.**
Xenium probe panel design

**Supplementary Table 3.**
Xenium experiment QC metrics

**Supplementary Table 4.**
Spatially varying genes and enriched ligands along gland-pit axis

**Supplementary Table 5.**
Raw counts and metadata bulk RNAseq growth factor screen

**Supplementary Table 6.**
Ionocyte-enriched DE genes with binary tissue overlap and cross-dataset enrichment annotations

**Supplementary Table 7.**
EEC-subtype DEGs and TF enrichment bins

**Supplementary Table 8.**
TF Modules inferred by Pando

**Supplementary Table 9.**
Raw counts and metadata bulk RNAseq ASCL1 and NEUROG3 overexpression

**Supplementary Table 10.**
Differential expression analysis of NEUROG3 and ASCL1 overexpression
